# SAVANA: reliable analysis of somatic structural variants and copy number aberrations in clinical samples using long-read sequencing

**DOI:** 10.1101/2024.07.25.604944

**Authors:** Hillary Elrick, Carolin M. Sauer, Jose Espejo Valle-Inclan, Katherine Trevers, Melanie Tanguy, Sonia Zumalave, Solange De Noon, Francesc Muyas, Rita Cascão, Angela Afonso, Fernanda Amary, Roberto Tirabosco, Adam Giess, Timothy Freeman, Alona Sosinsky, Katherine Piculell, David T. Miller, Claudia C. Faria, Greg Elgar, Adrienne M. Flanagan, Isidro Cortes-Ciriano

## Abstract

Accurate detection of somatic structural variants (SVs) and copy number aberrations (SCNAs) is critical to inform the diagnosis and treatment of human cancers. Here, we describe SAVANA, a computationally efficient algorithm designed for the joint analysis of somatic SVs, SCNAs, tumour purity and ploidy using long-read sequencing data. SAVANA relies on machine learning to distinguish true somatic SVs from artefacts and provide prediction errors for individual SVs. Using high-depth Illumina and nanopore whole-genome sequencing data for 99 human tumours and matched normal samples, we establish best practices for benchmarking SV detection algorithms across the entire genome in an unbiased and data-driven manner using simulated and sequencing replicates of tumour and matched normal samples. SAVANA shows significantly higher sensitivity, and 9- and 59-times higher specificity than the second and third-best performing algorithms, yielding orders of magnitude fewer false positives in comparison to existing long-read sequencing tools across various clonality levels, genomic regions, SV types and SV sizes. In addition, SAVANA harnesses long-range phasing information to detect somatic SVs and SCNAs at single-haplotype resolution. SVs reported by SAVANA are highly consistent with those detected using short-read sequencing, including complex events causing oncogene amplification and tumour suppressor gene inactivation. In summary, SAVANA enables the application of long-read sequencing to detect SVs and SCNAs reliably in clinical samples.

## Main

Cancer genomes are shaped by diverse forms of structural variants (SVs), ranging from simple deletions, inversions and amplifications, to complex patterns involving hundreds to thousands of SVs across multiple chromosomes^1–4^. SVs underpin most cancer driver mutations, including oncogene amplification and inactivation of tumour suppressor genes, and underlie the somatic copy number aberrations (SCNAs) frequently observed in cancer cells^1,2,4^. To date, the study of cancer genomes has largely relied on short-read whole-genome sequencing (WGS) on the dominant Illumina platform, which generates short, highly accurate reads of 100-150bp. However, detection of SVs using short reads is limited because breakpoints falling in repetitive regions cannot be unambiguously aligned to the human genome, and the strong GC bias of Illumina sequencing limits power for SV detection^5,6^. As a result, the landscape of SVs in cancer genomes remains partially mapped.

Long-read sequencing methods, such as the platforms commercialised by Oxford Nanopore Technologies (ONT) and Pacific Biosciences (PacBio), permit continuous reading of individual DNA molecules up to megabases^7,8^. While initially limited by high per-sample cost, high DNA input requirements, low and non-uniform flow cell yield, and high sequencing error rates, long-read sequencing methods are now mature enough to enable the study of diverse types of genomic variants, including SVs, the complete characterization of genes systematically missed by short reads^9^, and gapless genome assembly^10^. The potential of long reads to characterise complex somatic SVs at single-haplotype resolution, such as viral integration events^11–13^ and complex rearrangements^14–20^, has catalysed interest in the application of ONT and PacBio sequencing to study structural variation processes in human cancers. Initial long-read studies of cancer cell lines and small sets of clinical samples reported thousands of somatic SVs only detectable in long-read data, hinting at a hidden landscape of somatic SVs recalcitrant to short-read sequencing^21^. However, a recent analysis revealed that most SVs underlying SCNAs larger than 10 kbp are detected by short-read sequencing, raising the question as to whether the higher rates of SVs detected in previous studies using long-read sequencing reflect true biological signal or false positive calls due to limited algorithmic performance.

Over the last few years, several algorithms specifically designed for the detection of SVs using long reads have been developed^20,22–28^. To date, assessment of algorithmic performance has primarily relied on SV truth sets encompassing small sets of tens SVs detected in cancer cell lines and validated experimentally^20,22–28^. However, using SV truth sets to benchmark SV detection methods is limited in multiple ways. First, SV truth sets are biased towards genomic regions amenable to experimental validation. This results in a bias towards excluding SVs in low-complexity regions, which are precisely the genomic elements intractable to short-read sequencing that could potentially be characterised more accurately using long reads. Second, the selection of candidate SVs for experimental validation is performed using existing SV detection algorithms, which might have their own biases (e.g., sensitivity for SV detection might be variable across SV classes or genomic regions). Third, SV truth sets are only reliable to evaluate the recall of algorithms, but not their specificity because, without additional experimental validation, it is not possible to determine whether SVs called outside the genomic regions interrogated experimentally are artefacts or true SVs. Finally, experimental validation of SVs is time-consuming and not easily scalable, which limits the size of SV truth sets, and thus, the statistical power for benchmarking algorithms. Therefore, due to the lack of both large-scale long-read data sets for human cancers and best practices for benchmarking somatic SV detection algorithms in an unbiased manner, the relative performance of existing algorithms and the extent to which long-read sequencing approaches can improve the characterization of SVs, and by extension SCNAs, as compared to short-read sequencing remains unclear.

To address these needs, we present SAVANA, a computationally efficient algorithm specifically designed to detect both somatic SVs and SCNAs, as well as infer tumour purity and ploidy, using long-read sequencing data, including the estimation of tumour purity and ploidy. Through benchmarking experiments of PCR-validated rearrangements and sequencing replicates applied to long-read data from cancer cell lines and clinical samples, we show that SAVANA significantly outperforms existing algorithms in both sensitivity and specificity across long-read sequencing platforms, genomic regions, clonality levels, SV types and SV sizes. Analysis of matched, high-depth nanopore and Illumina WGS data for a collection of 99 clinical samples spanning diverse cancer types, revealed that SAVANA detects most SVs and SCNAs detected by Illumina, while also discovering additional rearrangements which were not detectable by short-read sequencing. In addition, tumour purity and ploidy estimates computed using SAVANA strongly correlated with those estimated using a short-read WGS analysis pipeline suitable for delivering clinical tumour sequencing reports. SAVANA is compliant with VCF specifications to facilitate downstream analyses, is implemented in Python 3 and is available at https://github.com/cortes-ciriano-lab/savana.

## Results

### Overview of SAVANA

We developed SAVANA to detect both somatic SVs and SCNAs in long-read sequencing data, including the estimation of tumour purity and ploidy (**Fig. 1a and Methods**). In brief, SAVANA scans sequencing reads from a tumour and matched normal sample pair jointly to identify clusters of alignments supporting the same SV. Given that the length of long reads is heterogeneous and alignment accuracy at breakpoints depends on read length, the same SV (*e.g*., a deletion) might be supported by sequencing alignments either fully or partially spanning the breakpoints, which leads to different types of evidence, *i.e*., gapped alignments and split alignments, respectively. Therefore, alignments supporting the same SV type (including gapped and split alignments), mapping within a given distance to each other, and in the case of insertions also of the same size range, are considered to support the same putative SV (**Methods**). SAVANA also includes functionalities to detect single breakends, in which only one of the two genomic regions bridged by an SV can be unambiguously mapped to the reference genome, thus enabling the detection of SVs involving low complexity or repetitive regions, such as centromeres or retrotransposons, and insertions of sequences not included in the reference genome, such as viral insertions (**Methods**). In addition, SAVANA provides functionalities to assign sequencing reads supporting each breakpoint to haplotype blocks when the input sequencing reads are phased.

**Figure 1.**
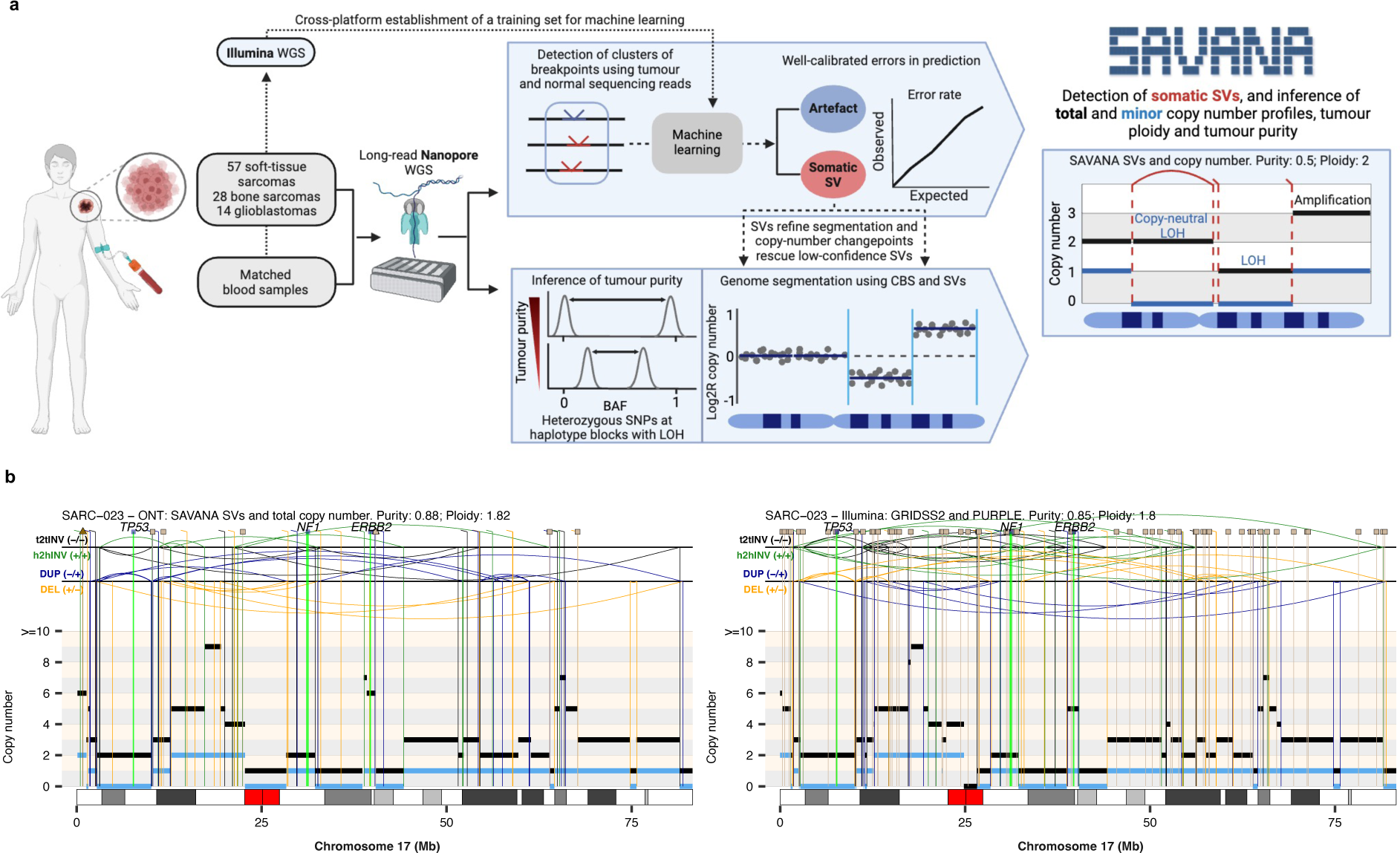
Overview of SAVANA. (a) Methodology for the analysis of somatic SVs, SCNAs, tumour purity and ploidy using SAVANA. (**b**) Example of a complex genomic rearrangement profile detected using SAVANA and matched short-read sequencing data. The total and minor allele copy-number data are represented in black and blue, respectively. DEL, deletion-like rearrangement; DUP, duplication-like rearrangement; h2hINV, head-to-head inversion; t2tINV, tail-to-tail inversion. Lines with a square at the top represent single breakends, and lines with arrowheads mark insertions. BAF: B-allele frequency; CBS: circular binary segmentation; LOH: loss of heterozygosity; Mb: megabase pairs; SV: structural variant; WGS: whole- genome sequencing.

A key innovation of SAVANA is the use of machine learning to distinguish somatic SVs from sequencing and mapping errors. Specifically, SAVANA encodes each breakpoint using a set of covariates related to the location, type and size of the event, orientation of reads at the breakpoint junction (*e.g.*, + for the upstream breakpoint of a deletion^1^), number of supporting alignments and depth of coverage (**Methods**). These features serve to predict whether a given cluster of SVs is consistent with either a true somatic SV or sequencing errors using a machine learning model trained on a large collection of SVs detected using matched long-read sequencing and short-read sequencing data (**Methods**). In this way, true SVs are distinguished from clusters of SVs arising from both recurrent and non-recurrent artefacts.

To detect SCNAs, SAVANA utilises somatic breakpoints and circular binary segmentation (CBS) to partition the genome into regions showing equal read depth (**Fig. 1a and Methods**). In addition, the intersection of somatic SVs and copy number changepoints is used to rescue SVs that did not have enough evidence to be called based on sequencing read alignment information only. Next, SAVANA infers the tumour purity by considering the mean B-allele frequency (BAF) values of heterozygous SNPs at regions with loss of heterozygosity (LOH). The key idea is that in a tumour without infiltration of non-neoplastic cells, the number of sequencing reads supporting a lost allele should be zero, and thus, the BAF values at regions with LOH should be 0 or 1. However, the infiltration of normal cells in the tumour results in a shift of the BAF values towards 0.5. The magnitude of this deviation is proportional to the degree of normal infiltration, thus allowing inference of tumour purity^29^. For this calculation, SAVANA considers sample-specific heterozygous SNPs mapping to the same copy number segment or haplotype block when the sequencing reads provided are phased (**Methods**). SAVANA uses the copy number information to rescue breakpoints that did not pass the thresholds for being called using read alignments if they support copy number changepoints, thus increasing sensitivity. Finally, SAVANA determines the ploidy of the tumour and the allele-specific copy number profile that best explains the observed sequencing read depth and BAF data given the estimated tumour purity (**Fig. 1b and Methods**).

### Establishing a training set for somatic SV detection using machine learning

We performed high-molecular weight DNA extraction on 99 tumour-normal pairs (57 diverse soft tissue sarcomas, 28 osteosarcomas and 14 glioblastomas), and sequenced the DNA using both nanopore WGS (median sequencing depth of 51x and 34x for tumour and normal samples, respectively) and Illumina WGS (118x and 41x for tumour and normal samples, respectively) (**Fig. 1b, Supplementary Fig. 1, Supplementary Table 1 and Methods**). We achieved a median N50 of 15 and 21kbp for tumour and normal nanopore sequencing reads, respectively. After stringent QC of the short and long-read data, including the removal of samples showing high rates of artefactual fold-back-like inversions (**Supplementary Fig. 2 and Methods**), 84 tumours were considered for further analysis (**Supplementary Table 1**).

To assemble a high-quality training set of true somatic SVs (true positives) and artefacts (true negatives), we used a clinical-grade short-read WGS data analysis pipeline^30^ to detect somatic SVs in the Illumina data (**Methods**). Next, we labelled SVs detected by SAVANA in the long-read data as true somatic SVs (*n*=52,464) if they were also detected in the short-read data, and as false positives otherwise (*n*=14,282,014; **Methods**). To prevent misclassifying true somatic SVs only detectable by long-read sequencing as false positives, we excluded from the training data SVs mapping to repetitive regions, blacklist regions^31^, and regions amplified to a high copy number in the tumour (>200 reads), as well as SVs overlapping with germline insertions detected in a panel of normals (**Methods**).

To assess the generalisation of our approach to distinguish true from false positive SVs, we used Random Forest (RF) classification to train leave-one-tumour-out models. Specifically, we classified the SVs detected in each tumour using an RF model trained on the SVs detected in all other tumours in the cohort. Overall, model performance was high, with a mean area under a receiver operating characteristic curve (AUC) of 0.98 (range 0.97-0.98). The most predictive covariates included the number of supporting reads and alignments in the tumour and matched normal sample, SV length, the number of unphased alignments supporting a breakpoint, the tumour allele fraction, and the count of normal reads clustered at the same locus supporting any breakpoint orientation (**Supplementary Fig. 3 and Supplementary Table 2**).

By default, SAVANA uses RF classification to identify somatic SVs. However, SAVANA also provides functionalities to assess the reliability of individual predictions and ensure high model performance for the minority class (true somatic SVs in this case) using Mondrian Conformal Prediction (MCP). While some SV detection algorithms provide quality scores^32^, these scores are not formally guaranteed to be well calibrated, since the fraction of erroneous predictions does not correlate with class probability scores. By contrast, MCP guarantees mathematically that the fraction of predictions for which the set of predicted labels does not include the true class is not higher than a user-defined tolerated error provided that the interchangeability assumption is satisfied, i.e., that the training and test examples are independently and identically distributed, which is a common requirement in predictive modelling. The maximum error rate is ensured for each category, including the minority class (somatic SVs), which is particularly important when modelling highly-imbalanced data sets^33,34^, as is the case here. In practice, MCP predicts four classes: true positive (somatic SV), false positive, *Null* and *Both*. The *Null* category indicates that at the chosen confidence level the model considers the SV to be too dissimilar to the somatic and false positive SVs in the training data to make a reliable prediction, indicating that the instance is an outlier (**Supplementary Fig. 4)**. By contrast, the *Both* category is assigned to instances that are similar to both the somatic SVs and false positives in the training data, and thus, cannot choose a single category. In practice, *Null* and *Both* predictions are considered to be erroneous and correct, respectively. Increasing the confidence level forces the model to reduce the number of *Null* predictions, which might cause an increase in SVs classified as *Both*^33^.

Overall, MCP provides a flexible and mathematically sound approach to assess the reliability of individual SV calls.

### SAVANA outperforms existing SV detection algorithms

Next, we sought to compare the performance of SAVANA against existing SV detection algorithms specifically developed for long-read data, including widely used methods originally designed for germline SV discovery and have been frequently used for the detection of somatic SVs (Sniffles2^28^, CuteSV^24^ and SVIM^23^), as well as algorithms developed to detect somatic SVs using matched tumour-normal data (NanomonSV^20^ and Severus^22^; **Methods**). The total number of SVs detected in each tumour varied by up to two orders of magnitude across SV callers (**Supplementary Figs. 5-6**). To assess whether the variable rates of somatic SVs reported by existing algorithms and SAVANA are the result of variable sensitivity or specificity, we first benchmarked algorithm performance using a truth set of 68 somatic SVs detected in the melanoma cell line COLO829 and validated using either PCR or capture-based technologies^35^ (**Methods**). To this aim, we analysed long-read PacBio and ONT WGS data from COLO829 and a matched normal cell line, COLO829-BL (**Methods**). Overall, SAVANA showed significantly higher recall and specificity than existing algorithms across sequencing platforms (PacBio and ONT) and flow cell versions (*P* < 0.0001, two-sided Wilcoxon test, **Supplementary Fig. 7**).

We next sought to establish best practices for benchmarking and quantifying the performance of somatic SV detection algorithms across the entire genome in an unbiased manner, thus overcoming the limitations of SV truth sets for benchmarking SV callers. Specifically, we propose the use of simulated sequencing replicates. The key point here is that true somatic SVs should be detected in all replicates, whereas false positive calls resulting from library preparation or sequencing errors should only be detected in one replicate. The use of replicates has been widely established as an efficient strategy to benchmark algorithms for the detection of point mutations and indels^36–38^. More broadly, both technical and biological replicates represent a cornerstone across various research domains to assess the reproducibility and robustness of experimental and computational results, including sequencing data analysis^37,39^. Yet, the use of replication to benchmark somatic SV detection algorithms is surprisingly lagging.

Here, we simulated replicates in silico by randomly splitting the sequencing reads from each tumour sample into two different BAM files and applied each SV detection algorithm on each replicate independently. This strategy allows to simulate sequencing replicates with the same sequencing depth, thus overcoming the impact that uneven flow cell yield across sequencing replicates could have on SV detection sensitivity. Next, we compared the sets of somatic SVs detected by each algorithm across replicates (**Fig. 2a**). We found that the number of somatic SVs detected in just one replicate varied by up to two orders of magnitude across algorithms (**Fig. 2b**). Consistent with its higher performance on the COLO829 data set, SAVANA showed a significantly higher concordance across replicates compared to other algorithms across diverse tumour types (*P* < 0.01, two-sided Wilcoxon test, **Fig. 2c and Methods**).

**Figure 2.**
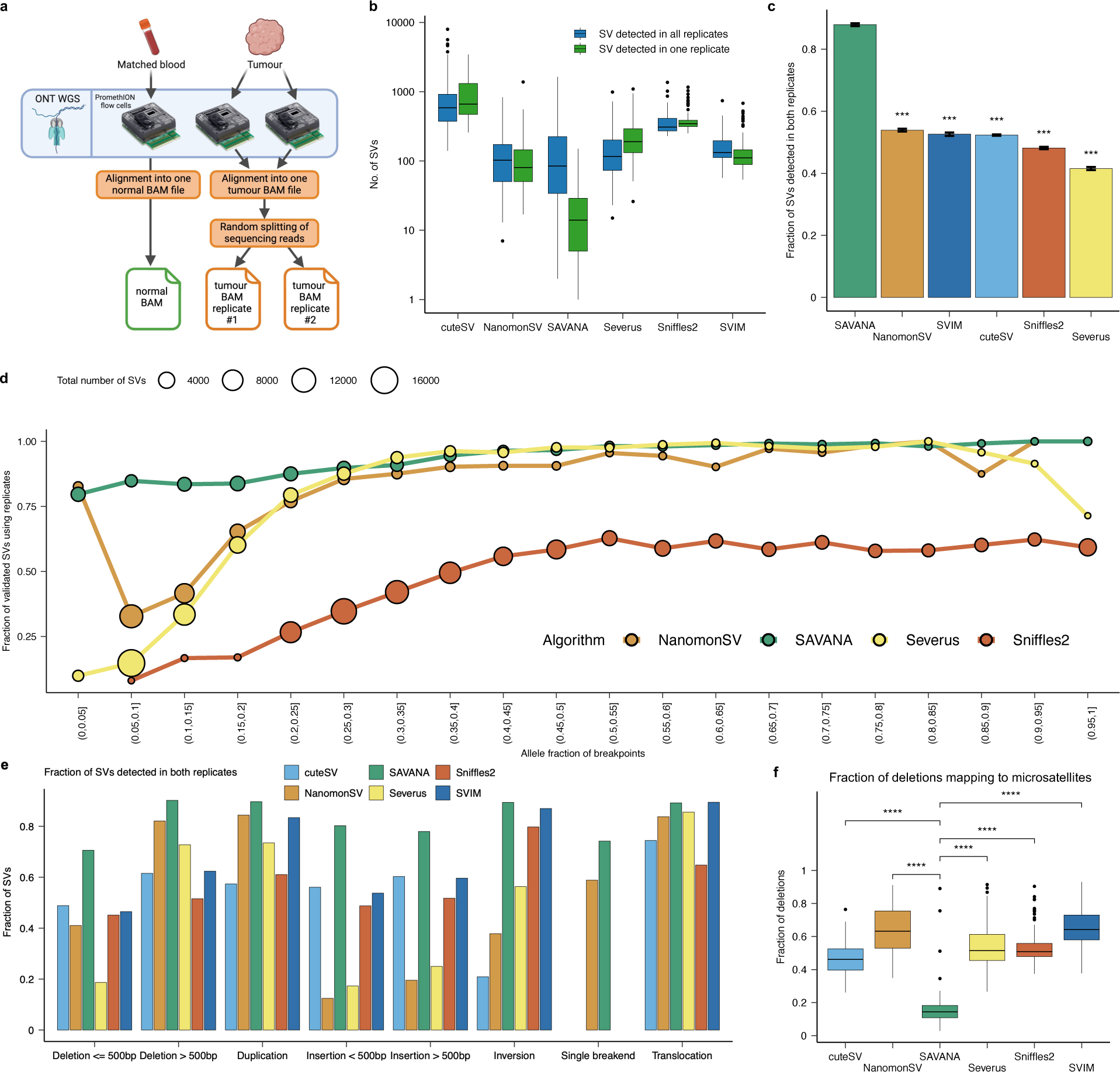
Benchmarking of SAVANA against existing algorithms using replicates. (**a**) Schematic representation of the replicate analysis strategy implemented to benchmark the performance of somatic SV detection algorithms. (**b**) Distribution of the number of somatic SVs detected by each algorithm stratified based on whether they were detected in one (green) or both (blue) replicates. (**c**) Comparison of the fraction of somatic SVs detected in both replicates by each caller. The bars report the result across all samples. The error bars report the 95% bootstrap confidence interval computed using 1000 bootstrap resamples. Significance with respect to SAVANA was assessed using the two-sided Student’s *t*-test (\*\*\**P* < 0.0001). (**d**) Fraction of somatic SVs detected in both replicates as a function of allele fraction. The size of the dots represents the number of somatic SVs in each group. Only algorithms that report the number of reads supporting each breakpoint or the allele fraction of breakpoints were included in this analysis. (**e**) Comparison of the fraction of somatic SVs detected in both replicates by each caller stratified by SV type. Each bar represents the fraction of replicated SVs across all samples. Panels **b-e** show the aggregated results for the 53 tumours with the highest sequencing depth. (**f**) Fraction of deletions mapping to microsatellite regions. Significance was assessed using the two-sided Wilcoxon’s rank test (\*\*\*\**P* < 0.00001).

A potential limitation of using replicates to benchmark mutation detection algorithms is that tumours often show high levels of intra-tumour genetic heterogeneity. As a result, true somatic SVs present at low allele frequency (AF) in a tumour might only be detected in one replicate if the number of sequencing reads in other replicates is not enough to meet the threshold for SV calling^38,39^. Thus, we next sought to investigate whether the variable number of SVs detected by different algorithms is the result of variable sensitivity for low-AF SVs. By analysing the fraction of SVs detected in both replicates as a function of AF (**Fig. 2d**) or the number of reads supporting each SV (**Supplementary Fig. 8**a), we found that SAVANA shows a higher and uniform concordance rate across SV clonality levels. In addition, SAVANA uniformly yielded the highest fraction of concordant SVs across tumours (**Supplementary Fig. 8**b), different classes of repeats (**Supplementary Fig. 8**c), SV types (**Fig. 2e**) and SV sizes (**Supplementary Fig. 9**). The higher concordance across replicates, especially in the low-AF range, indicates that SAVANA shows both higher specificity and sensitivity than existing methods.

We found that the relative number of SVs of each SV type was highly variable across algorithms, with cuteSV, Severus, Sniffles2 and SVIM detecting thousands of deletions and insertions (**Supplementary Fig. 10**). In addition, the concordance across replicates was highly variable across SV types, with insertions and deletions showing the lowest concordance rates (**Fig. 2f**). These results would be consistent with the higher error rate expected for ONT sequencing at low complexity regions, such as homopolymers^40,41^, suggesting that these SV types might contribute a large fraction of false positive calls. However, the number of deletions and insertions detected by these algorithms in one or two replicates was comparable. Because artefactual SVs mapping to genomic regions prone to sequencing or mapping errors might be detected in both replicates, we next sought to investigate whether the high rates of insertions and deletions reported by existing algorithms represent true biological events or artefacts. Analysis of the genomic distribution of SVs revealed that most of the insertions and deletions detected by these algorithms, especially those smaller than 500bp, mapped to microsatellites (**Fig. 2f and Supplementary Figs. 11-12**). By contrast, the rate of insertions and deletions detected by SAVANA at microsatellite loci was significantly lower (*P* < 0.0001, two-sided Wilcoxon test, **Supplementary Figs. 11-12**). In fact, the total number of insertions and deletions at microsatellites detected by the other algorithms was two to three orders of magnitude higher than SAVANA for all tumour types analysed (**Supplementary Fig. 12**). High rates of insertions and deletions at microsatellites, a phenotype known as microsatellite instability (MSI), are associated with inactivation of the mismatch repair pathway^42,43^. However, the rates of MSI in the tumours analysed in this study are very low^42^, and we did not detect inactivation of MMR genes or MSI using matched Illumina WGS data for the tumours analysed here. Therefore, taken together, these results indicate that the high rates of insertions and deletions at microsatellite loci reported by existing SV detection algorithms are most likely false positive calls resulting from sequencing or mapping errors. By contrast, the rates and types of SVs detected by SAVANA are consistent with tumour biology.

### Using replicates of matched germline controls to quantify false positive rates

Next, to quantify the specificity of SV detection algorithms, we harnessed sequencing and simulated replicates of matched germline controls (**Fig. 3a**). Specifically, we first ran each algorithm on sequencing replicates of COLO829BL using one replicate as the tumour and the other one as the matched germline control. The key idea here is that an algorithm with optimal specificity should not detect any somatic SVs in this setting, thus, any somatic SVs detected represent false positive SV calls. On the COLO829BL data set, SAVANA showed the lowest false positive rate, reporting 7 false positive SVs (**Fig. 3a**). In comparison, the other algorithms showed 9x (NanomonSV: 66 false positives), 59x (SVIM: 411), 69x (486 Severus), 277x (Sniffles2: 1940) and 391x (CuteSV: 2737) higher false positive rates (**Fig. 3a**). To investigate the specificity of SV detection algorithms in clinical samples, we next performed the same analysis using simulated replicates for the blood WGS data generated as matched germline controls for the tumour samples sequenced in this study. To this end, we randomly split the sequencing reads from the matched normal sample into two BAM files (**Fig. 3b**). Next, each algorithm was run using one replicate as if it was the tumour sample and the other as the matched normal. Consistent with the results for the COLO829BL replicates, SAVANA showed a significantly lower false positive rate as compared to other algorithms, which showed tens to thousands of false positives per sample (*P* < 0.0001, two-sided Wilcoxon test, **Fig. 3c**). Notably, the order of magnitude difference between existing methods and SAVANA is comparable for both the number of false positives (**Fig. 3b**) and the number of somatic SVs detected (**Supplementary Figs. 5-6**), thus suggesting that the variable number of somatic SVs detected in tumours is driven by the low specificity of existing algorithms. Investigation of rearrangement profiles revealed that somatic SVs detected by SAVANA largely agree with Illumina SV calls and that breakpoints map to copy number changepoints both detected using short-read and long-read data (**Fig. 4a-b and Supplementary** Figs. 13-18). In contrast, existing algorithms detect high rates of SVs, especially insertions and deletions, which are not supported by somatic copy number changepoints, while also missing SVs detected by SAVANA and present in Illumina WGS data (**Fig. 4c-f and Supplementary Figs. 13-18**).

**Figure 3.**
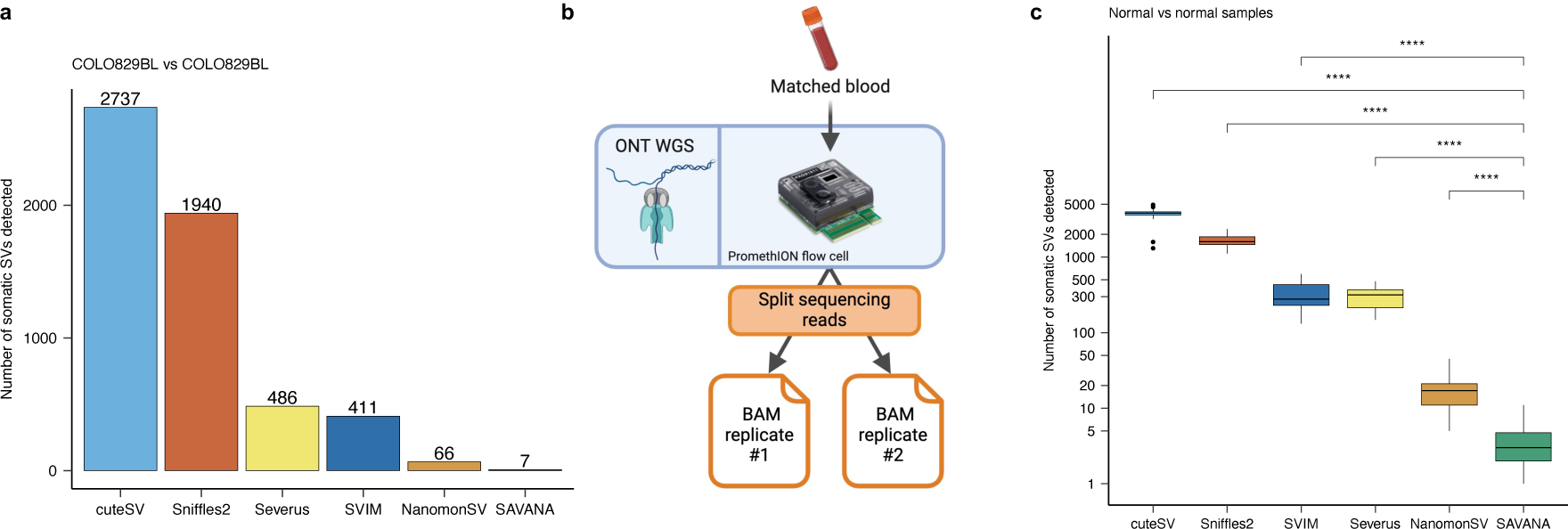
Benchmarking the specificity of SAVANA against existing algorithms using sequencing replicates of matched germline controls. (**a**) Number of somatic SVs detected in the COLO829BL cell line when running the algorithms benchmarked using a normal replicate as the tumour sample. The number on top of each bar indicates the number of false positive calls for each algorithm. (**b**) Schematic representation of the replicate analysis strategy implemented to quantify the false positive rate of somatic SV detection algorithms. (**c**) Distribution of false positive SV calls detected when running the SV detection algorithms benchmarked using replicates of whole-blood normal samples generated in silico by splitting sequencing reads randomly into two BAM files. Each dot represents one donor. Significance in (**c**) was assessed using the two-sided Wilcoxon’s rank test (\*\*\*\**P* < 0.00001).

**Figure 4.**
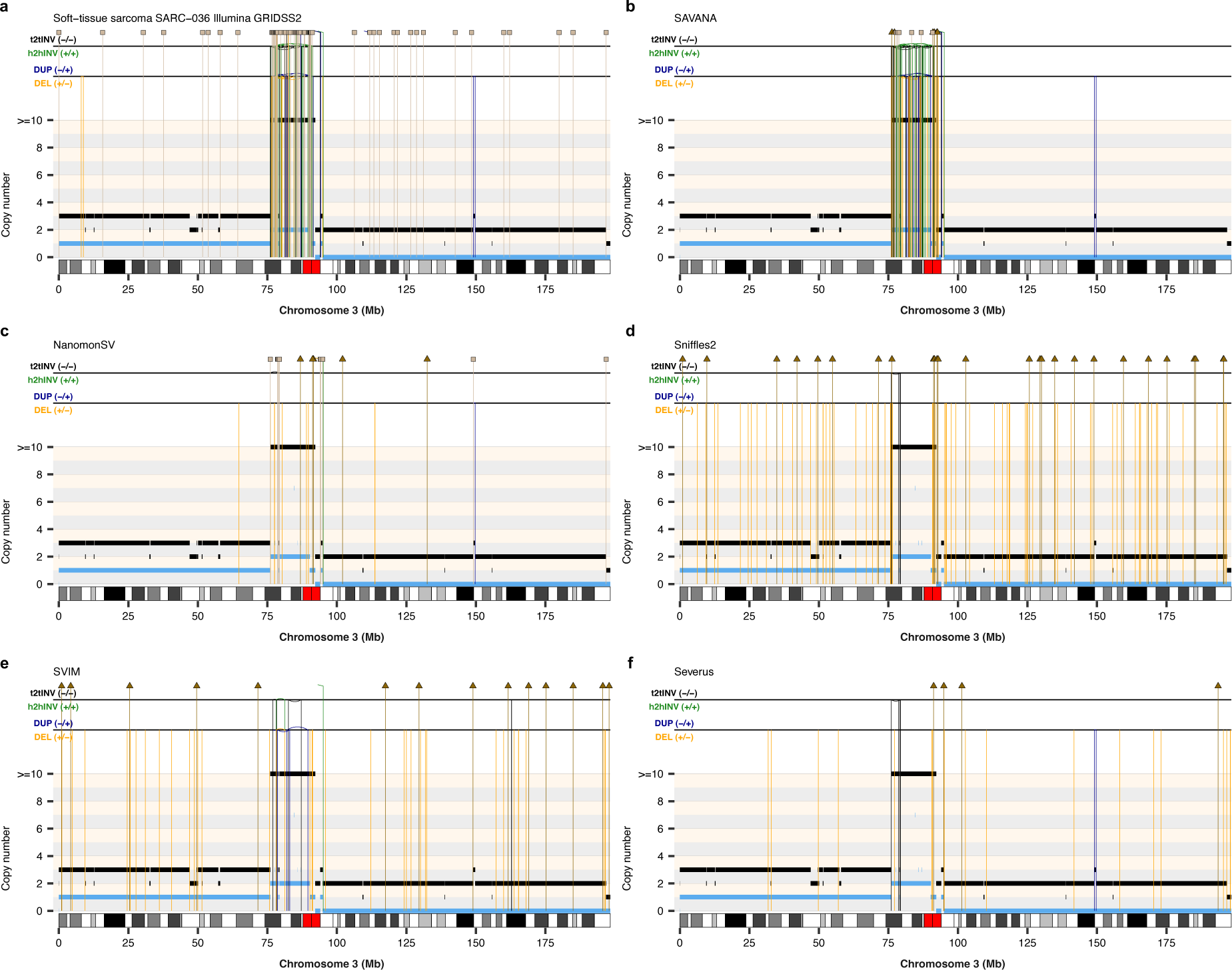
Benchmarking of SV detection algorithms. (**a**) Somatic SVs and copy number profiles detected using GRIDSS2 and PURPLE in whole- genome short-read sequencing data. Somatic SVs detected in matched long-read nanopore whole-genome sequencing data using SAVANA (**b**), NanomonSV (**c**), Sniffles2 (**d**), SVIM (**e**) and Severus (**f**). The copy number profiles shown in **a-f** were calculated using PURPLE and the short-read sequencing data. The total and minor allele copy-number data in **a-f** are represented in black and blue, respectively. DEL, deletion-like rearrangement; DUP, duplication-like rearrangement; h2hINV, head-to-head inversion; t2tINV, tail-to-tail inversion. Lines with a square at the top represent single breakends, and lines with arrowheads mark insertions.

Together, these results demonstrate that SAVANA consistently shows both significantly higher specificity and sensitivity than existing methods across diverse SV types and clonality levels, and further, that the high rate of somatic SVs detected by existing algorithms is explained by their low specificity rather than by higher sensitivity. Moreover, SAVANA offers significantly faster runtimes than existing algorithms (*P* < 0.0001, two-sided Wilcoxon test, **Supplementary Fig. 19**).

### SAVANA detects cancer driver SVs and SCNAs with high accuracy in clinical samples

Next, we set out to investigate the differences between long and short-read sequencing for somatic SV detection. On average, 85% of the SVs detected in the Illumina data by SAVANA were also present in the long-read data (**Fig. 5a-c, Supplementary Fig. 20 and Methods**). In contrast, the recall rate for other algorithms was significantly lower (*P* < 0.001; two-sided Wilcoxon test, **Fig. 5a-b**). Furthermore, using existing algorithms, no more than 40% of the SVs detected in the long-read sequencing was detected in the Illumina data (*P* < 0.0001; two-sided Wilcoxon test, **Fig. 5b and Supplementary Fig. 21**). These low recall rates are consistent with the lower concordance across replicates observed for existing algorithms (**Fig. 2**).

**Figure 5.**
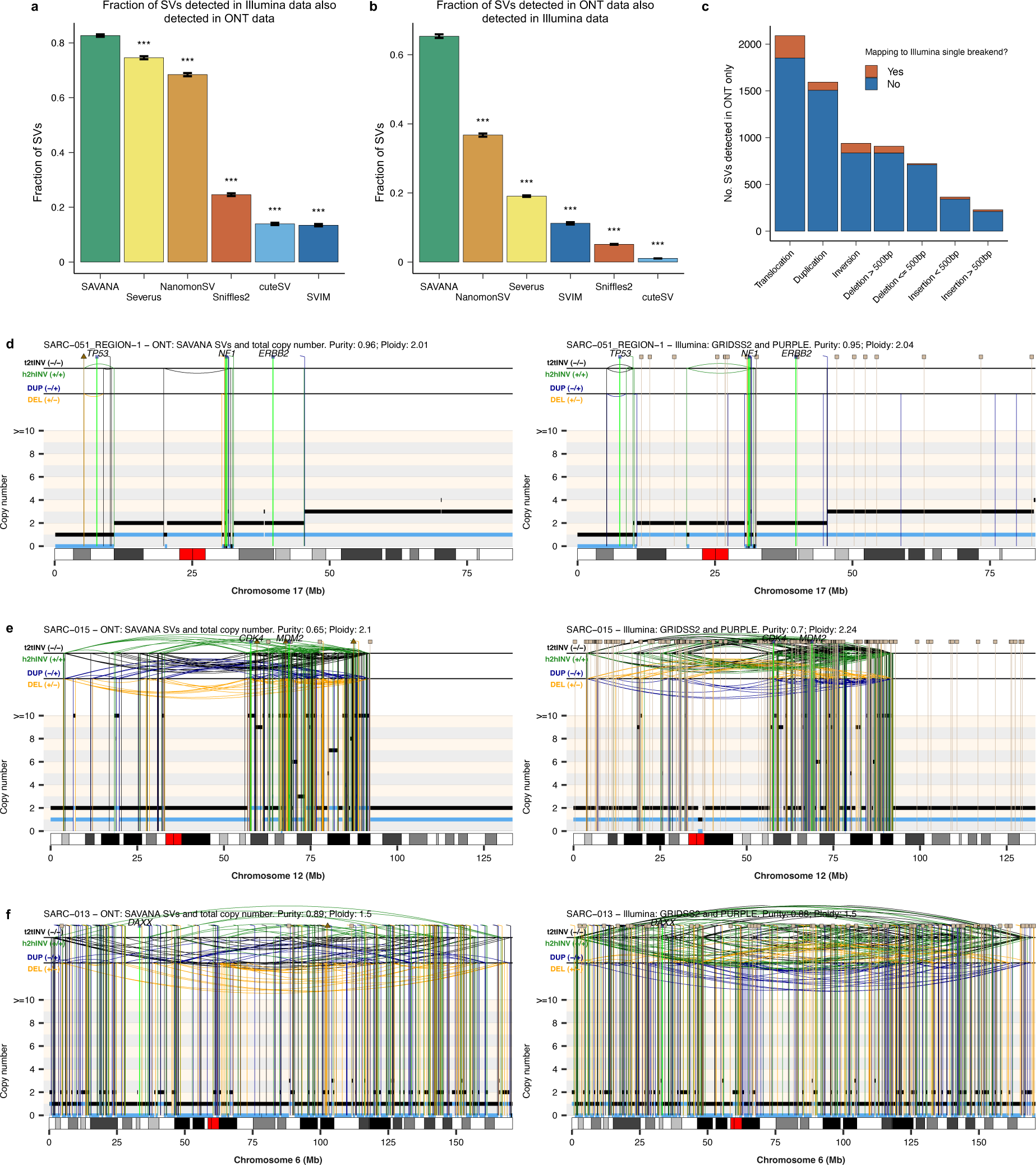
Comparison between short and long-read data for the analysis of SVs and SCNA. (**a**) Fraction of somatic SVs detected in Illumina WGS data using GRIDSS and PURPLE that are also detected in ONT data by the algorithms benchmarked. The fractions shown were computed by aggregating the somatic SV calls detected in all tumours in the cohort. (**b**) Fraction of somatic SVs detected in ONT WGS data by each of the algorithms benchmarked that were also present in Illumina WGS data. Significance in **a-b** was assessed using the two-sided Wilcoxon’s rank test (\*\*\**P* < 0.001). (**c**) Total number of SVs across the cohort detected in ONT data using SAVANA stratified based on whether at least one breakpoint mapped to a single breakend detected using GRIDSS2. **(d-f**) Examples of somatic SVs and SCNAs detected in long-read nanopore whole-genome sequencing data using SAVANA (left) and in Illumina WGS data using GRIDSS2 and PURPLE (right). The total and minor allele copy-number data are represented in black and blue, respectively. DEL, deletion-like rearrangement; DUP, duplication-like rearrangement; h2hINV, head-to-head inversion; t2tINV, tail-to-tail inversion. Lines with a square at the top represent single breakends, and lines with arrowheads mark insertions.

In addition to somatic SV calling, SAVANA integrates read depth and B-allele frequency information to infer tumour purity, ploidy and allele-specific copy number. To evaluate the performance of SAVANA for SCNA analysis, we first compared the SCNAs computed by SAVANA against the combination of PURPLE and GRIDSS2 applied to Illumina WGS data. These analyses showed that the tumour purity and ploidy values estimated by SAVANA and PURPLE/GRIDSS2 are strongly correlated (Pearson’s correlation coefficient of 0.97 and 0.9, respectively, **Fig. 6**). These results are noteworthy given the higher than two-fold difference in median sequencing depth between the Illumina and nanopore data sets (118x vs 51x, respectively), and that, as opposed to PURPLE, SAVANA does not use the AF values of somatic point mutations to score purity and ploidy combinations.

**Figure 6.**
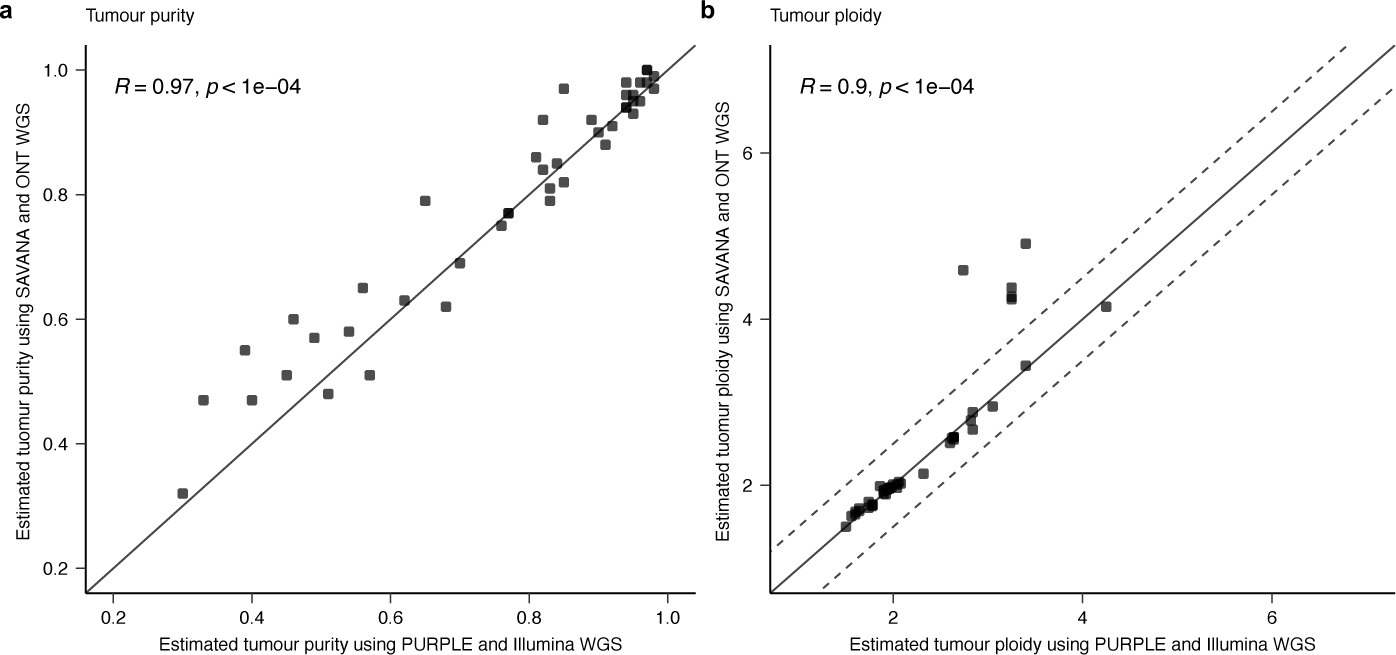
Comparison of the tumour purity and ploidy estimates computed using SAVANA and ONT WGS data against PURPLE and Illumina WGS data. R: Pearson’s correlation coefficient. For this analysis, we only considered the 44 tumours with region-matched nanopore and Illumina WGS data.

We next set out to investigate genomic rearrangements mediating the inactivation of tumour suppressor genes and the amplification of oncogenes. SAVANA detected diverse types of complex rearrangements encompassing SVs and SCNAs leading to the loss of tumour suppressor genes, such as chromothripsis events causing the inactivation of *CDKN2A, NF1, TP53* or *RB1* **(Fig. 5d-f and Supplementary** Figs. 22-31). In addition, SAVANA detected complex genomic rearrangements mediating oncogene amplification across various tumour types, such as amplification of *CDK4*, *MDM2, CCNE1* and *MYC* in osteosarcomas, and *MYC* and *EGFR* in glioblastoma^1^.

A major advantage of long-read sequencing is that long-range phasing information provides the opportunity to phase SVs and other types of mutations^16^. To enable haplotype-aware analysis of SVs, SAVANA provides functionalities to assign SVs to individual haplotypes if the sequencing reads provided as input are phased using germline SNPs. Subsequently, the haplotype-resolved SV calls generated by SAVANA permit the application of genome assembly tools to reconstruct derivative chromosomes resulting from complex SVs in clinical samples (**Supplementary Fig. 32**).

Overall, most SVs and SCNAs detectable by state-of-the-art short-read sequencing analysis pipelines can be detected in long-read data using SAVANA, including the inactivation of tumour suppressor genes and the amplification of oncogenes.

## Discussion

Here, we show that SAVANA enables the integrative analysis of somatic SVs, SCNAs, tumour purity and ploidy using long-read sequencing data. Through the analysis of a large collection of matched Illumina and ONT WGS data for clinical and non-clinical samples, we show that SAVANA shows significantly higher sensitivity and specificity than existing methods across a dynamic range of clonality levels, SV sizes and SV types. This is critical for the analysis of clinical samples given that tumour purity is often low (down to 20%) for multiple tumour types. In practice, this means that driver mutations might be missed if sensitivity for low-AF SVs is low even when sequencing is performed to high depth. Notably, the specificity of SAVANA is 9- and 59-times higher specificity than the second and third-best performing algorithms, respectively, while also showing higher sensitivity. Consequently, SAVANA enables enhanced detection and analysis of SVs and more robust interpretation of the underlying tumour biology. Our genome-wide benchmarking analysis of SV detection algorithms has revealed that existing methods show low specificity for somatic SV detection, which results in hundreds to thousands of false positive somatic SV calls per tumour. Thus, our results strongly suggest that the high rates of somatic SVs previously reported for cancer genomes using long-read sequencing do not correspond to true biological signal. To facilitate reliable and consistent comparison of algorithms in the future, we establish best practices for benchmarking somatic SV detection methods using replicates that allow quantification of the sensitivity and specificity of algorithms across the entire genome in an unbiased manner.

The large scale of our tumour sequencing data set and the high accuracy of SAVANA allowed us to compare the relative performance of long and short-read sequencing for somatic SV detection. SAVANA detects most of the SVs detected by Illumina sequencing, including complex events that mediate the inactivation of tumour suppressors or amplification of oncogenes. Nevertheless, we note that our results are limited by the read lengths obtained for the samples included in this study, which might be too short to characterise somatic SVs in long stretches of low-complexity sequences, such as centromeres. Thus, ultra-long read sequencing data sets will be needed to fully chart the landscape of somatic SVs in cancer genomes.

SAVANA utilises machine learning to distinguish artefacts from true somatic SVs. Because the diversity of the training data determines the performance of any machine learning model, it is essential to control for prediction confidence to increase the reliability and adoption of algorithms based on predictive modelling. In other words, it is essential to provide quantitative metrics to answer the following question: what level of performance will users achieve on new data and how reliable will the results be? To this end, SAVANA provides functionalities to assess the applicability domain of the models using MCP^44,45^. MCP enables the detection of instances falling outside the applicability domain of the model and provides well-calibrated metrics to quantify prediction confidence, which have proven transformative in diverse applications, such as drug discovery^33^, spatial transcriptomics^46^ and computational histopathology^47^. In this study, we have illustrated how MCP can be used to control the error rate for the somatic SV category. However, MCP could be further extended to control the error rate for e.g. different SV types or genomic regions, or integrated with other algorithms, such as neural networks^48^. Finally, the fact that SAVANA uses non-identifiable predictive features for training MCP models opens avenues for improving model performance by integrating genomics data in a federated manner. For example, MCP model training could be distributed without the need for sharing raw sequencing data, thus facilitating data integration while also ensuring compliance with local data privacy regulations and not comprising proprietary data, as shown in drug discovery applications^49^.

Overall, the high performance of SAVANA for the joint analysis of SVs and SCNAs in clinical samples will facilitate the reliable application of long-read sequencing to study structural variation processes in cancer genomes and the detection of clinically relevant alterations in human tumour samples.

## Methods

### SAVANA algorithm

SAVANA consists of the following steps:

#### - Discovery of putative breakpoints

SAVANA uses aligned reads from tumour and normal BAM files to identify breakpoints. Specifically, SAVANA splits the genome into non-overlapping bins of 100 million bp by default, and the alignments from each bin are examined in parallel. Thus, the analysis of large chromosomes is distributed across multiple processes and small chromosomes are analysed in a single process. Primary and supplementary alignments with a mapping quality over the minimum threshold of 5 are considered for further analysis. To ensure that all potential germline SVs are identified and used to discount germline SVs as somatic, a more lenient mapping quality filter is applied to the normal reads. Specifically, the minimum mapping quality threshold is lowered by a fraction of 0.5 for alignments originating from the normal sample.

For each sequencing read, SAVANA parses supplementary alignments (SA) from the SA tag and creates breakpoint objects storing the location, orientation, phasing, and read name information. If the single breakend option is enabled, supplementary alignments with low mapping quality (default < 5) are considered unmapped and their sequence and location are recorded as potential single breakends.

SAVANA parses the CIGAR string for both primary and supplementary alignments, tracking deletion and insertion operations greater than the minimum SV length (30bp by default), and inserted bases in the case of insertion operations. Large deletions can be interspersed with short regions of mapped sequence. To account for this phenomenon, adjacent deletions are merged and reported as a single putative event when interspersed by mapped sequences shorter than 30bp. Soft-clip operations in the CIGAR string greater than 1000bp (by default) without a corresponding supplementary alignment are also tracked at this stage, as well as the soft-clipped bases. To more accurately remove germline SVs from the tumour SV set, the minimum SV length to track an insertion or deletion is reduced by a fraction of 0.20 for the normal BAM file. This step prevents reporting as somatic those germline SVs falling below the minimum length cutoff used to analyse the tumour reads.

In addition, while iterating through aligned reads to identify putative breakpoints, a coverage array is built to track the count of overlapping primary and supplementary alignments, separated by their haplotype. For each haplotype (including none for unphased reads), an array of zeros is created for each contig, with an element for each bin across the genome. The value of the array position corresponding to the bin overlapped by an alignment is then incremented by 1 for the corresponding haplotype and contig array. This array is later used to annotate the depth of sequencing at breakpoints by accessing the number of overlapping reads from each haplotype, as well as the sum of these. The bin size is defaulted to five bases, which can be increased by the user to reduce memory usage and run time.

Putative SVs are recorded using breakend notation following the variant call format (VCF) specifications. In addition, SVs are annotated using the notation scheme established by the Pan-Cancer Analysis of Whole Genomes (PCAWG) project^1^: + - for deletion-like SVs, - + for duplication-like SVs, + + and - - for inversion-like SVs, and INS for insertions. Putative deletions derived from the CIGAR string are reported as ‘+-’ to ensure that supplementary and gapped alignments supporting the same deletion event are grouped together.

#### - Clustering of putative breakpoints

Once information about individual putative breakpoints is collected, adjacent breakpoints are clustered together if they fall within a clustering window. By default, this clustering window is 10bp for all breakpoint types except insertions. We use a clustering window of 250bp for insertions because we frequently observed that insertions typically have more variance in their mapping location than other breakpoint types. Reads supporting each breakpoint are split into clusters based on the contig of mate breakpoints. A final clustering step is performed on the reads supporting each mate breakpoint using a larger clustering window (default of 50bp) to prevent supporting reads from being incorrectly split into multiple events. Insertion events are clustered by their insert size to ensure that grouped reads support insertions of comparable size. Finally, grouped events are split by their respective breakpoint notation, and their supporting information and location in the genome are stored in a dictionary. Single breakends are reported if they are the only type of event present at a given breakpoint cluster. This is required because the detection of single breakends adjacent to other types of events might indicate that some of the sequencing reads in the cluster were long enough to permit complete reconstruction of the underlying SV, whereas other reads only allowed reliable mapping of one side of the new adjacency, thus resulting in single breakends. For example, an insertion of a repetitive sequence might be fully spanned by some sequencing reads, thus allowing complete reconstruction of the event. However, other reads might span only a small fraction of the inserted sequence, thus hampering unambiguous mapping of the inserted sequence, which would be detected as a single breakend.

#### *-* Filtering of false positives using machine learning

Breakpoints are encoded using 38 features, including the average MAPQ of supporting reads, the mean, standard deviation, and median SV length, the standard deviation of the start and end mapping locations of supporting reads, etc. (see **Supplementary Table 2** for a description of all breakpoint features computed). Sequencing depth in the tumour and normal sample before, at, and after each breakpoint is also computed by summing the coverage array from the first base of the contig to the breakpoint start coordinate as described above. If the input BAMs provided are phased, the haplotype of supporting reads and their phase sets are also reported in the VCF output by SAVANA.

To differentiate clusters of noise or sequencing artefacts from true somatic or germline events, we trained a Random Forest (RF) classifier using Scikit-Learn^50^. To this aim, we annotated all breakpoints detected by SAVANA in the long read data sets as true somatic SVs if they were also detected as somatic SVs in matched Illumina data or as noise otherwise. To establish a high-confidence set of SVs for model training, we excluded breakpoints which (1) were not found in Illumina but were however supported by at least six tumour reads in the ONT data; (2) mapped within 500bp of a repetitive element reported by RepeatMasker^51^; (3) mapped within a microsatellite; (4) mapped to an ENCODE blacklist region^31^; (5) mapped to regions with high read depth in the tumour sample (>200 reads); and (6) intersected with the location of four or more putative insertions from across the cohort with at least five supporting reads in either a tumour or normal sample.

Features encoding breakpoints (**Supplementary Table 2**) are read in from raw VCF files and transformed into a Pandas dataframe^52^. Breakpoints are then split into training and test sets with a 4:1 ratio. We used default parameter values to train the RF models except for the number of trees and the maximum depth. The optimal values for these parameters were determined using cross validation and grid search using a range of 100 to 1000 for the number of trees and 10 to 25 for the maximum depth. To detect somatic SVs on each tumour, we trained an RF model using the labelled SVs from all other tumours to ensure that no data from the tumour to be analysed was used during model training.

Once the RF model is trained, the raw SAVANA VCF file is labelled with an additional CLASS variable in the INFO column that denotes the model prediction. The predictions of the model are used to set the FILTER column to ‘PASS’ if the breakpoint is predicted to be somatic or ‘LIKELY_NOISE’ otherwise. The model predictions are vetted by additional thresholds on the minimum allele fraction and number of supporting reads, and their filter column is updated to “LOW_SUPPORT” or “LOW_AF” if they do not meet minimum thresholds of 0.01 for allele fraction and 3 supporting reads by default. Conversely, breakpoints classified as false by the model with at least 7 tumour supporting reads, a minimum tumour allele fraction of 0.10, a mean read mapping quality over 15 and no clusters of breakpoints of any type mapping to the same location in the normal sample, are rescued and their FILTER column updated to ‘PASS’.

#### - Reliability estimation of SV calls using conformal prediction

Mondrian Conformal Predictors were implemented as previously described^33,34^. In brief, the training data was randomly split into two sets: 70% of the datapoints were used to train an RF model, which was applied on the remainder. The fraction of trees voting each class was used as the nonconformity measure. We generated one list of nonconformity measures for each class, which were then used to compute *P* values using a predefined confidence level. The *P* values were then used to assign breakpoints the one of the following categories: somatic, likely noise, *Both* or *Null*.

### Detection of somatic copy number aberrations

SAVANA provides functionalities for the detection of total and allele-specific copy number aberrations and the estimation of tumour purity and ploidy.

#### - Genome-wide read depth computation

First, SAVANA generates non-overlapping bins of 10kbp incorporating SAVANA breakpoints (if provided) across the reference genome. Each bin is annotated with the percentage of non-N bases and its overlap with blacklisted regions when provided. Next, SAVANA counts the number of tumour and normal sequencing reads mapping to each bin across the genome, excluding bins with more than 5% overlap with blacklisted regions and bins encompassing more than 75% non-N bases in the reference genome. Secondary and supplementary read alignments, as well as reads with a mapping quality < 5, are not considered for this analysis. Bins that do not have any read counts are filtered and removed. Binned tumour read counts are then normalised using normal counts from the matched normal sample while accounting for differences in read depth between tumour and matched normal BAM files. Subsequently, binned read counts are log2 transformed providing relative log2 copy number ratio values (log2R). Next, single point outliers in the log2R read counts are smoothened, as previously described^53^, to reduce noise prior to segmentation of the log2R data. Subsequently, log2R read counts are segmented using circular binary segmentation (CBS), an algorithm for finding changepoints in sequential data^54^, as follows. First, copy number changepoints are identified using 1,000 permutations setting the threshold for statistical significance to 0.05. Changepoints are then validated using 1,000 permutations and a threshold for statistical significance of 0.01. To handle potential oversegmentation, resulting genomic segments are subsequently merged if their log2R difference is smaller than the 20th percentile of the distribution defined by the log2R difference between all pairs of segments.

#### - Inference of tumour purity

To infer tumour purity, SAVANA computes the B-allele frequency (BAF) for all germline heterozygous SNPs in the genome. Next, SAVANA searches for haplotype blocks showing evidence of LOH. To this end, SAVANA computes the bimodality coefficient of the BAF distribution for each haplotype block larger than 500kbp and with at least 10 heterozygous SNPs. Haplotype blocks with read depths larger than twice the mean read depth are excluded, as potentially amplified or gained regions would bias the BAF distribution. Subsequently, the median BAF values are computed for the top ten haplotype blocks with the highest bimodality coefficient value for each of the parental alleles (BAFA and BAFB), and tumour purity is estimated using the following equation:

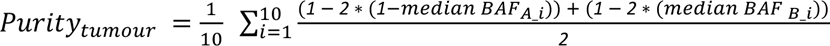

#### - Inference of tumour ploidy

Once the tumour purity (Puritytumour) is estimated, SAVANA infers tumour ploidy using grid search, as previously described^55^, encompassing a purity search space of {0.01* ∣ * ∈ [Puritytumour - 0.05 .. Puritytumour + 0.05]} range, and ploidy search space of {0.01* ∣ * ∈ [1.5 .. 5]} range. Each purity-ploidy combination is subsequently assessed for its goodness-of-fit by computing the root mean squared deviation (RMSD) or mean absolute deviation (MAD) weighted by the segment size between the inferred absolute copy number and the nearest integer values. In addition, for each purity-ploidy combination, SAVANA evaluates whether the proportion of copy number states fitted to zero is < 0.1, the proportion of copy number segments that are considered close to the next nearest integer is > 0.5, and the copy number change step size between the two most frequent copy number states is < 2. Purity-ploidy combinations which do not pass these criteria are discarded. The purity-ploidy combination with the best goodness-of-fit (i.e. lowest RMSD or MAD value) is finally used to compute the absolute copy number for each segment using the observed log2R data as follows^55^:

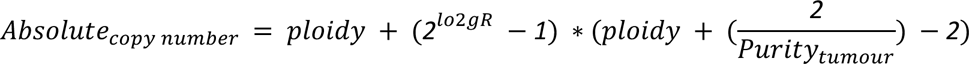

#### - Allele-specific copy number fitting

Finally, the optional tumour purity and ploidy values estimated in the previous step are used to compute the allele specific copy number profile for the tumour as previously described^56^.

### Selection of human sarcoma samples and matched germline controls

Fresh-frozen bone and soft-tissue sarcoma samples were obtained from patients consented and enrolled in both the Genomics England 100,000 Genomes Project (G100k) as well as the Royal National Orthopaedic Hospital (RNOH) Biobank, satellite of the UCL/UCLH Biobank for Health and Disease (REC reference 20/YH/0088). Patients did not receive financial compensation for donating samples. Surplus tumour tissue from resection and/or biopsy samples were collected and frozen as part of routine clinical practice. Matched blood samples were used for germline sequencing. Processing of tissue samples for pathology review and molecular analyses was performed at the RNOH for the 100,000 Genomes Project, founded by England’s National Health Service in 2012, as previously described^57^. Fresh-frozen tissue sections [haematoxylin and eosin (H&E); 5 μm] were used to guide the selection of the most viable areas for each tumour specimen in terms of lack of necrosis and tumour cellularity. DNA was extracted from matched tumour and blood samples using established protocols and in accordance with the 100,000 Genomes Project guidelines. DNA was sent for centralised library preparation and sequencing at the Illumina Laboratory Services (ILS) in Cambridge, UK^58^. Sequencing was performed as part of the 100K Genomes Project. The DNA from tumour and normal samples was sequenced to an average depth of 116 (median 118) and 42 (median 36), respectively. Dual consented somatic and germline genomic data were shared with RNOH and EMBL-EBI. Data linked to local clinical data (no NHS Digital or NHS England data) were shared by Genomics England with RNOH and EMBL-EBI. No analysis was undertaken in the Genomics England Research Environment or National Genomic Research Library.

### Selection of glioblastoma tumour samples and matched germline controls

Glioblastoma and matched blood samples were collected in the Neurosurgery Department at Centro Hospitalar Universitário Lisboa Norte (CHULN) and stored less than 1 h after surgery at Biobanco-iMM CAML (Lisbon Academic Medical Center, Lisbon, Portugal). Ethical approval was obtained from the Ethics Committee of CHULN (Ref. N° 367/18). Written informed consent was obtained from all patients prior to study participation in accordance with the European and National Ethical Regulation (law 12/2005). Patients did not receive financial compensation for donating samples.

### DNA extraction and processing for nanopore sequencing

Tumour genomic DNA was extracted from tissue with the Nanobind tissue kit (PacBio SKU 102-302- 100) and Germline DNA from blood with the Nanobind CBB kit (PacBio, SKU 102-301-900). After extraction, the genomic DNA (gDNA) was homogenised by 3-10 passes of needle shearing (26G) and 1h of incubation at 50°C. All samples were quantified on a Qubit fluorometer (Invitrogen, Q33226) with the Qubit BR dsDNA assay (Thermo Fisher Scientific, Q32853) and manual volume checks were performed. The DNA size distributions were assessed at each relevant step by capillary pulse-field electrophoresis with the FemtoPulse system (Agilent, M5330AA and FP-1002-0275). 4-10 μg of gDNA were either fragmented to a size of 10-50kb with the Megaruptor 3 (Diagenode, E07010001 and E07010003) or fragmented to a size of 15-25kb with a gTUBE (Covaris, 520079) centrifuged at 1,500rcf. All samples were depleted of short DNA fragments (less than 10kb long) with the SRE kit (Circulomics, SS-100-101-01, now PacBio) or the SRE XS kit (Circulomics SS-100-121-01, now PacBio) when sample availability or concentration was limiting. To generate matched Illumina and nanopore tumour WGS data, DNA aliquots were tumour matched. Cases for which multi-region nanopore and Illumina tumour sequencing was performed, DNA aliquots were tumour and region matched.

### Library preparation and nanopore whole-genome sequencing

Sequencing libraries were generated from 600ng up to 1.8μg of gDNA with 1 library prepared for germline samples and 2 libraries prepared for tumour samples with the SQK-LSK110 or SQK-LSK114kits (Oxford Nanopore Technology, ONT), according to manufacturer’s recommendations with minor modifications. Briefly, the samples were end-repaired by adding 2 μl NEBNext FFPE DNA Repair Mix (NEB, M6630) and 3 μl NEBNext Ultra II End Prep Enzyme Mix (NEB, E7546), incubated for 10 min at room temperature followed by 10 min at 65 °C, then cleaned up with 1 × AMPure XP beads (Beckman Coulter, A63880) and eluted in 60 μl of Elution Buffer. The end-repaired DNA was ligated with 5 μl Adapter Mix (ONT, SQK-LSK110) using 8 μl NEBNext Quick T4 DNA ligase (NEB, E6056) at 21°C for up to 1h. The adapter-ligated DNA was cleaned up by adding a 0.4 × volume of AMPure XP beads. Sequencing libraries were quantified using the average peak size of the samples determined after sample preparation on a Femto Pulse system. 20fmol of the obtained sequencing libraries were loaded onto R9.4 flow cell (ONT), or 10fmol of libraries were loaded onto R10 flow cells and sequenced on a PromethION48 (ONT). The libraries were stored overnight in the fridge. After 24h, sequencing runs were paused and a DNAse treatment or nuclease flush (ONT, WSH-003) was performed and 20fmol of the libraries were re-loaded on the flow cells.

### Nanopore sequencing data analysis

Base-calling was performed using the high accuracy model of Guppy-4.0.11 and a qscore filter of 7. Sequencing reads were aligned to GRCh38 using either minimap2 (version 2.24)(68) with parameters “-ax map-ont-MD”. QC statistics and plots were generated using cramino^59^ version 0.14.5.

### Analysis of artefactual fold-back-like inversions

Fold-back-like inversion artefacts were detected by the presence of reads characterised by having two nearly identical alignments in forward and reverse orientations. As a result, the first alignment starts where the second alignment ends, and both alignments are of equal length (**Supplementary Fig. 2**). To estimate the number of fold-back-like inversion artefacts per sample, we analysed all aligned reads per sample and classified them as a fold-back-like inversion artefact when the following criteria were met: (1) the read had exactly one primary and one supplementary alignment, both with a minimum mapping quality of 20; (2) the primary and supplementary alignments overlapped but were mapped in opposite orientations; and (3) the distance between the alignment positions corresponding to the start and end of the read were less than 150bp apart from each other. To calculate the rate of fold- back-like inversion artefacts per sample, we determined the total number of reads by counting those reads with a primary alignment and a minimum mapping quality of 20. Samples with more than 5 million artefactual fold-back-like inversions were not considered for further analysis. The code used for this analysis can be accessed at https://github.com/cortes-ciriano-lab/ont_fb-inv_artefacts.

### Detection of SVs using existing algorithms

Sniffles2 (version 2.2.0), cuteSV (version 2.1.0), SVIM (version 1.4.2), Severus (version 1.0), and NanomonSV (version 0.5.0) were used for benchmarking. Germline callers (Sniffles2, cuteSV, and SVIM) were run separately on tumour and normal BAM files, all with a minimum SV length of 32bp. Sniffles2 was run with ‘--output-rnames’. cuteSV was run with recommended parameters: ‘-- max_cluster_bias_INS 100’, ‘--diff_ratio_merging_INS 0.3’, ‘--max_cluster_bias_DEL 100’, and ‘-- diff_ratio_merging_DEL 0.3’. SVIM was run in ‘alignment’ mode as pre-aligned BAM files were used. To subtract germline calls from somatic, coordinates of germline SVs for each tool were extracted from their resulting germline VCF. The start and end coordinates were extended by 1000bp and saved to a BED file. A bedtools^60^ subtraction of this file from the tumour VCF was performed, requiring a fraction of at least 0.01 of the tumour SV to be overlapped by the extended germline SV coordinates for it to be removed. Only tumour SVs with three or more supporting reads were considered. However, all germline SVs were considered. NanomonSV was run with the recommended, ‘--use_racon’ flag, ‘- -single_bnd’, as well as a control panel provided by the tool developers to reduce false positives (available at: https://zenodo.org/records/7017953). Severus was run using haplotyped BAM files, as recommended by the tool. In all cases, SVs mapping to alternative contigs, chromosome M, unplaced contigs, and the Epstein-Barr virus were removed. SVs detected in Illumina and ONT data were considered to support the same event if the underlying breakpoints mapped within 100bp of each other.

### Benchmarking SV detection algorithms

The breakpoint coordinates for the SVs in the COLO829 SV truth set from^60^ were lifted over to GRCh38. The SV truth set was downloaded from http://zenodo.org/records/4716169#.YL4yTJozYUE (truthset_somaticSVs_COLO829_hg38lifted.vcf), which was filtered to contain only breakpoints that were detected by the authors in ONT WGS data. The VCF file was curated to report insertions in a single entry.

SAVANA includes a module, termed evaluate, to compare VCF files. Specifically, the evaluate module compares breakends and reports them as matched when they are within an overlap buffer (100bp by default). The VCF files generated by the germline SV detection algorithms benchmarked were compared to the COLO829 somatic SV truth set VCF. Specifically, we assessed whether breakends in the truth set mapped within 100bp of the breakends reported by the algorithms benchmarked. These results were then used to compute the precision, recall and F-measure for each algorithm. The evaluate module in SAVANA accounts for both breakend format VCFs, where each breakend of an SV has a line in the VCF, and for VCF formats where both breakends are reported in the same line. We used the same criteria to benchmark the performance of SV detection methods using replicates.

### Assembly of derivative chromosomes at single-haplotype resolution using SAVANA SVs

For each phase set defined by Whatshap^61^, SV supporting reads identified by SAVANA were used as input to generate an assembly using wtdbg2^62^ and subsequently polished using Racon^63^.

### Processing of short-read whole-genome sequencing data

Sequencing reads were mapped to the GRCh38 build of the human reference genome using BWA-MEM^64^ version 0.7.17-r1188. Aligned reads were processed following the Genome Analysis Toolkit (GATK, version 4.1.8.0) Best Practices workflow to remove duplicates and recalibrate base quality scores^65^. NGSCheckMate was utilised with default options to verify that sequencing data from tumour-normal pairs were properly matched^66^. Somatic SVs were detected using GRIDSS^66^ (v2.12.0, https://github.com/PapenfussLab/gridss), which we ran using default parameter values. B-allele frequency (BAF) information for heterozygous SNP sites was collected using AMBER (v3.5) and read depth ratios for the reference and alternate alleles were computed using COBALT (v1.11). BAF and read depth ratios for heterozygous SNPs were used as input to PURPLE^67^ (v2.54) to estimate the purity, ploidy, and somatic copy number aberrations of the tumour samples. Microsatellite instability was assessed using PURPLE. AMBER, COBALT, and PURPLE are developed by the Hartwig Medical Foundation and are freely available on GitHub: https://github.com/hartwigmedical/hmftools. The implementation of the WGS data analysis pipeline used in this study is available at: https://github.com/cortes-ciriano-lab/osteosarcoma_evolution.

### Statistical analysis and visualisation

All statistical analyses were performed using R version 4.1.3. The level of significance for all statistical analyses was set at 0.05. No statistical method was used to predetermine sample size. Rearrangement and copy number profiles were visualised using the R package ReConPlot^68^ v0.1. Structural variants were read into R data frames for visualisation purposes using the R package StructuralVariantAnnotation^69^.

## Code availability

The code for SAVANA is available on GitHub at https://github.com/cortes-ciriano-lab/savana

## Data availability

WGS data from the participants enrolled in the 100,000 Genomes Project can be accessed via Genomics England Limited following the procedure described at: https://www.genomicsengland.co.uk/about-gecip/joining-research-community/. In brief, applicants from registered institutions can apply to join one of the Genomics England Research Networks, and then register a project. Access to the Genomics England Research Environment is then granted after completing online training. The short and long-read sequencing data from the glioblastoma samples are available under controlled access at EGA under the accession number EGAD00001012101. Data access can be granted via the EGA for a defined time period after successful completion of a data access agreement provided by the WTSI CGP Data access committee. ONT sequencing data for the COLO829 and COLO829BL cell lines^35^ using R9.4 MinION/GridION flow cells were downloaded from the European Nucleotide Archive (ENA; project ID PRJEB27698). The nanopore WGS data generated by Oxford Nanopore Technologies (Oxford, UK) for the cell lines COLO829 and COLO829BL using the Ligation Sequencing Kit v14 were downloaded from the Amazon Web Services S3 bucket s3://ont-open-data/colo_2023.04/. Finally, PacBio HiFi sequencing data for COLO829 and COLO829BL generated using the Revio system were downloaded from https://downloads.pacbcloud.com/public/revio/2023Q2/COLO829/.

## Author contributions

A.M.F. and I.C.-C. designed, supervised, administered, and obtained funding for the study. H.E. implemented the SAVANA functionalities for the detection of structural variants with input from C.M.S., J.E.V.-I., S.Z., F.M. and I.C.-C. C.M.S. and I.C.-C. developed the SAVANA functionalities for somatic copy number analysis with input from H.E. H.E., C.M.S., J.E.V.-I., S.Z., and I.C.-C. performed analyses and generated the figures, with input from A.M.F. H.E., J.E.V.-I., S.Z., A.G., and I.C.-C. performed nanopore sequencing data analysis. S.D.N, F.A., R.T. and A.M.F. performed pathology review of sarcoma samples. C.C.F., R.C., and A.A. collected and processed the glioblastoma and matched blood samples. M.T. and G.E. performed nanopore sequencing. K.T., M.T., A.G., T.F., K.P., D.T.M., A.S., G.E., A.M.F and I.C.-C. provided technical support. H.E., C.M.S., and I.C.-C. wrote the manuscript with input from all authors. All authors read and approved the final version of the manuscript.

## Acknowledgements

H.E., C.M.S., J.E.V.-I., S.Z., F.M. and I.C.-C. thank EMBL for funding. I.C.-C. thanks The Wellcome Trust for funding. C.M.S. acknowledges support and funding from the Marie Skłodowska-Curie grant 101106070. This project was supported by research grants from the Sarcoma Foundation of America (SFA 20-05), the Connective Tissue Oncology Society (Basic Science Sarcoma Research Award) and NF Research Initiative at Boston Children’s Hospital awarded to A.M.F. and I.C.-C. Provision of patients’ samples from the Royal National Orthopaedic Hospital (RNOH) was made possible through the Royal National Orthopaedic Hospital Pathology Department and the Research and Development Department, The Tom Prince Trust, The Rosetrees Trust, Skeletal Cancer Trust, Sarcoma UK, The Bone Cancer Research Trust and The Pathological Society of Great Britain and Ireland, over the last two decades. We also acknowledge support to A.M.F from the National Institute for Health Research, UCLH Biomedical Research Centre, and the UCL Experimental Cancer Centre. SDN is funded jointly by the Pathological Society of Great Britain and Ireland and the Jean Shanks Foundation. This research was made possible through access to data in the National Genomic Research Library, which is managed by Genomics England Limited (a wholly owned company of the Department of Health and Social Care). The National Genomic Research Library holds data provided by patients and collected by the NHS as part of their care and data collected as part of their participation in research. The National Genomic Research Library is funded by the National Institute for Health Research and NHS England. The Wellcome Trust, Cancer Research UK and the Medical Research Council have also funded research infrastructure. R.C., A.A. and C.C.F. acknowledge the support of Associação David Vaz, Bolsa João Lobo Antunes – GAPIC (iMM/FMUL) and Millennium bcp. All authors thank the computational resources provided by the European Bioinformatics Institute (EMBL-EBI). The authors acknowledge the Biobanco-iMM CAML, which enabled the collection, processing, and storage of tumour and blood samples from glioblastoma patients. The authors thank Dr. Emma McCargow and Dr. Hélène Louis dit Picard for technical and administrative support. We thank the patients and their families for their participation in this study. We thank all members of the Flanagan and Cortes-Ciriano laboratories for numerous discussions and feedback. Some figures were generated using BioRender.com.

---

**Supplementary Figure 1.**
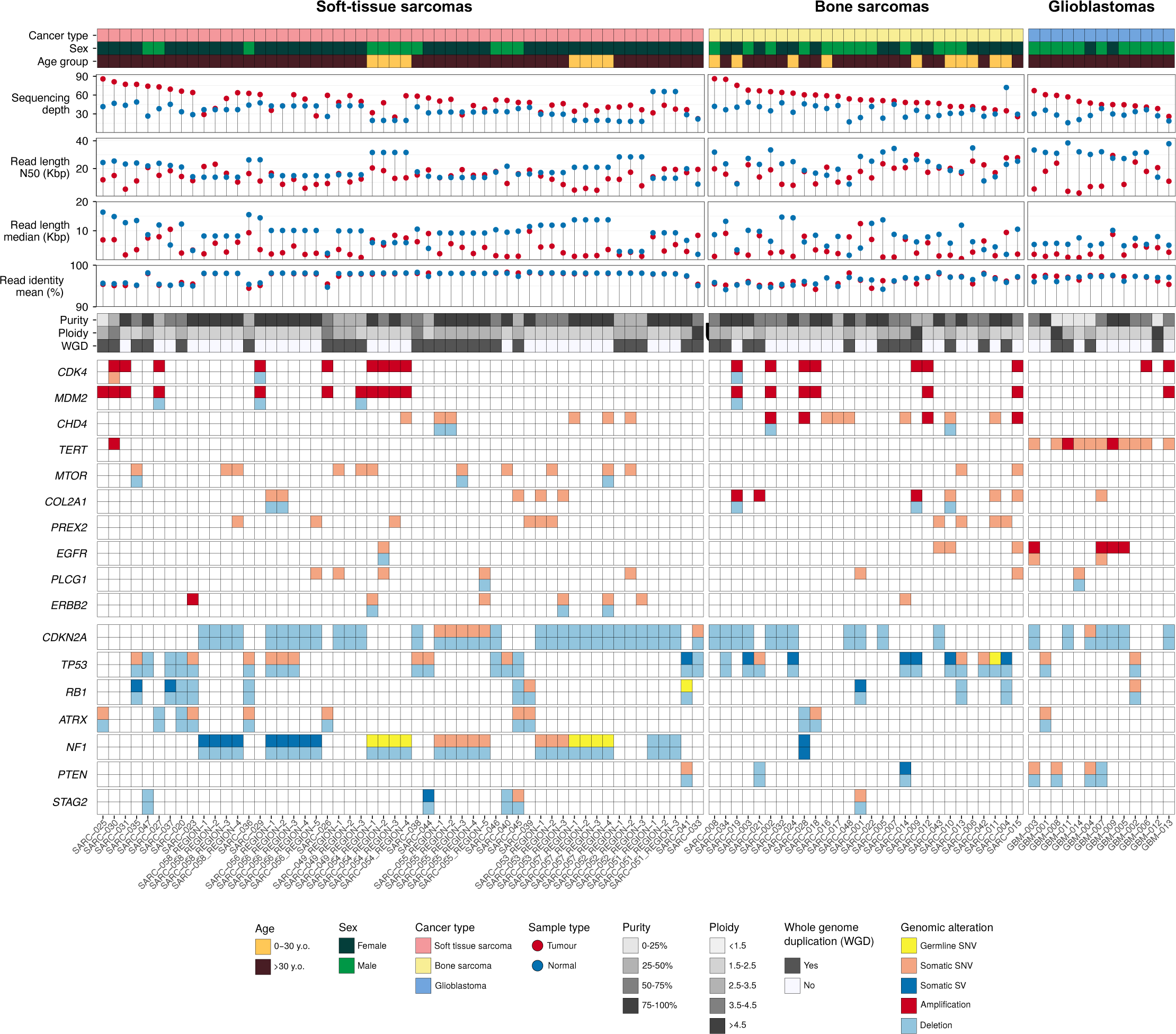
Overview of the tumour samples analysed in this study. Genomic and clinical landscape of the tumour samples analysed in this study using matched nanopore and Illumina WGS. Clinical information, histopathologic features, biallelic mutations and amplification of oncogenes are shown. The lollipop plots show the nanopore sequencing run results for the tumour and matched whole-blood samples. For *TERT*, only activating hotspot promoter mutations (c.-124C>T and c.-146C>T) were considered. Kbp: kilobase pairs; WGD: whole-genome doubling.

**Supplementary Figure 2.**
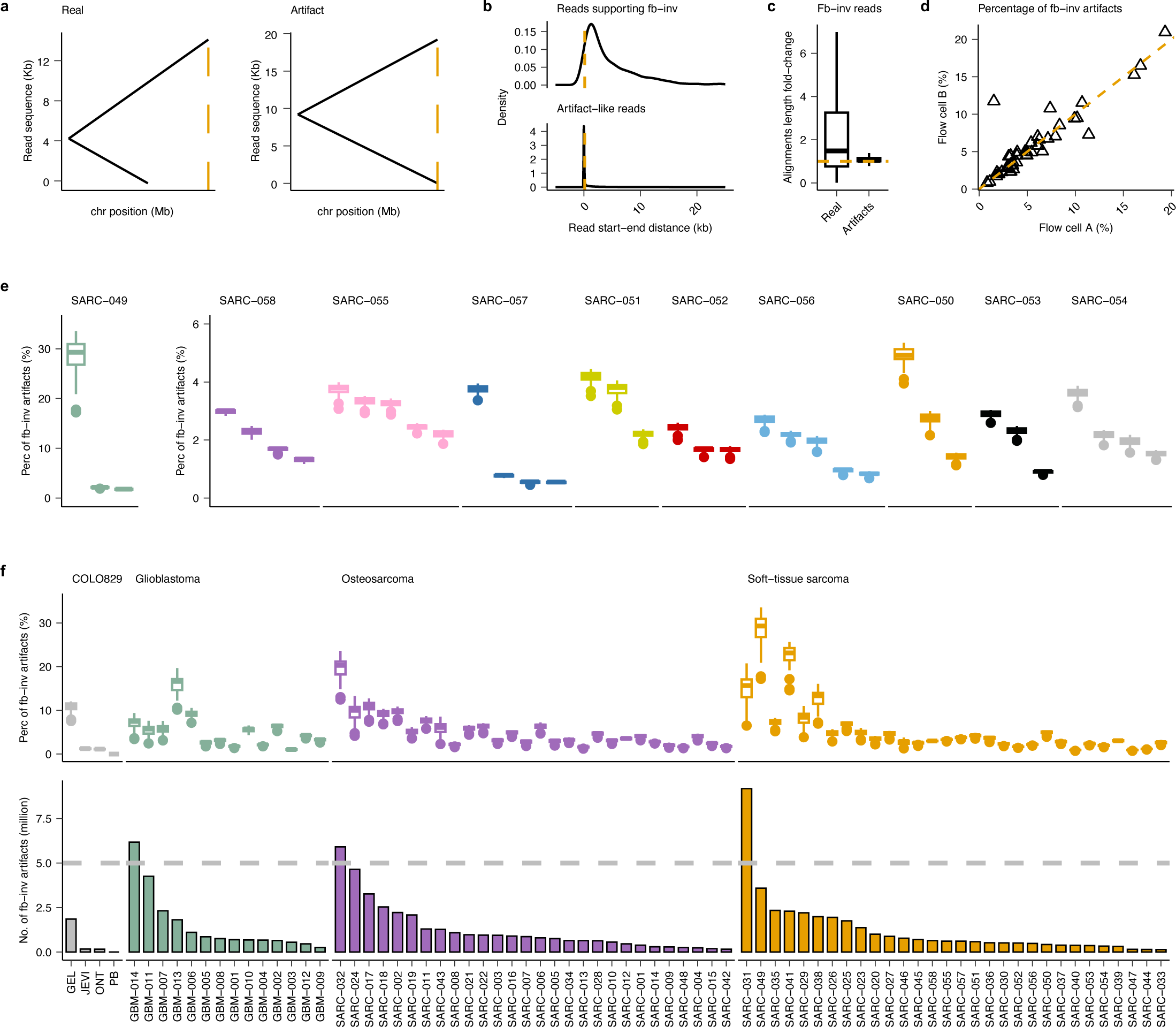
Analysis of fold-back-like inversion artifacts. (**a**) Scheme of a read alignment supporting a real fold-back inversion (left) and a fold-back-like artifact (right). The yellow dashed bar indicates the end alignment position in each case. (**b**) Distribution of the distance between the read start and end alignment positions. Although a uniform distribution is expected for genuine events, a peak is observed around 0, indicating the presence of artifact events. A threshold of 150bp WAS selected to classify these artefact reads. (**c**) Boxplot showing the fold-change of the forward and reverse alignment lengths. For reads supporting artifactual fold-back-like inversions, the forward and reverse alignment lengths are equal, resulting in a fold-change clustering around one. (**d**) Scatter plot showing the correlation of the percentage of artifacts between flow cell replicates. Dashed yellow line corresponds to the diagonal. (**e**) Boxplot showing the percentage of artifacts per chromosome across different tumour regions and patients. The differences are greater among regions than chromosomes, further supporting the artifactual origin. (**f**) Percentage of artifacts per chromosome (upper panel) and the total number of artifacts per tumour (lower panel) across the study cohort. For visualization purposes, the tumour region with the highest rate of artifacts was selected as the representative sample for each donor. The dashed grey line indicates the threshold of 5 million artifacts. tumours GBM-014, SARC-031 and SARC-032 were discarded according to this criterion. Fb-inv: fold-back inversion; Perc: percentage.

**Supplementary Figure. 3.**
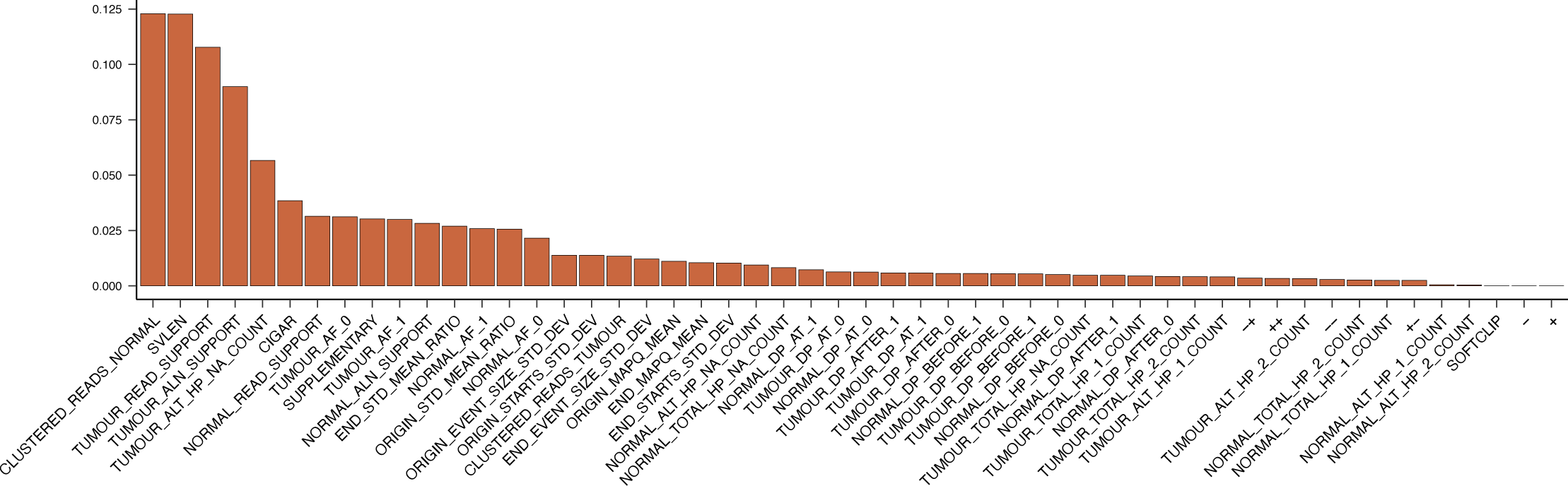
Mean decrease in accuracy (feature importance) obtained for each variable used in the Random Forest model to distinguish sequencing artefacts from true somatic SVs. A description for each feature is provided in Supplementary Table 2.

**Supplementary Figure 4.**
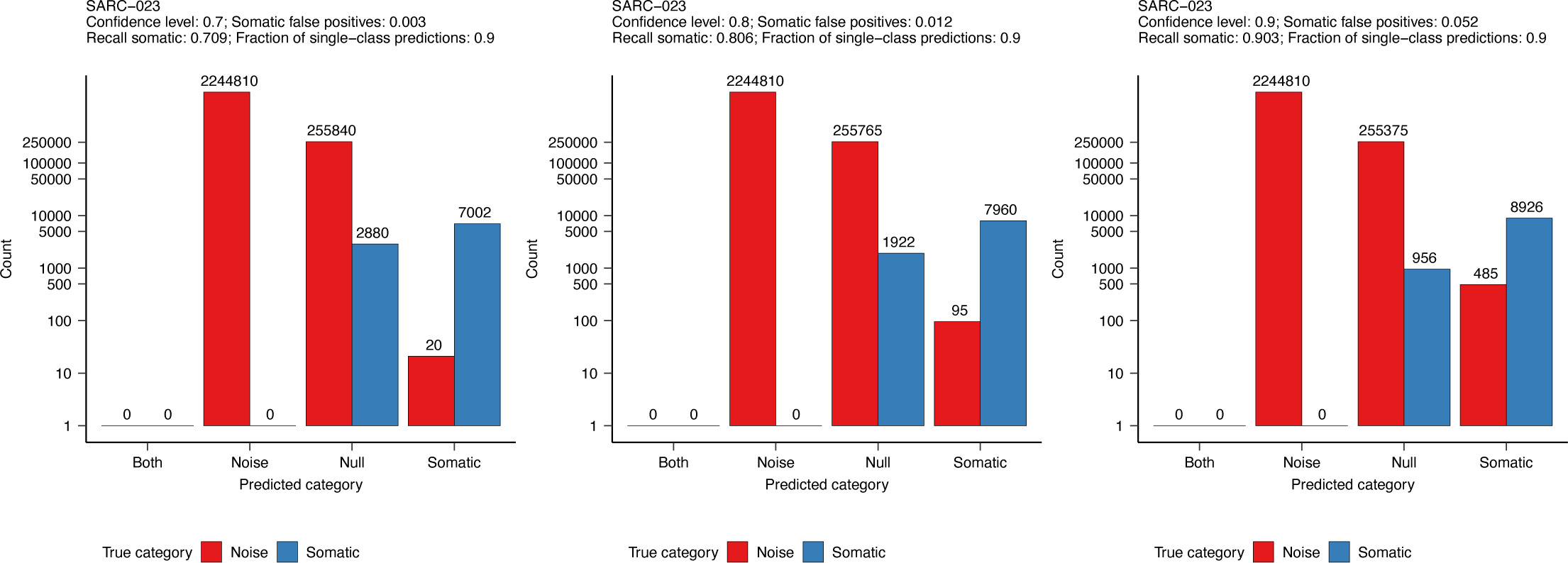
Analysis of the predicted categories using Mondrian Conformal Prediction. The bars show the number of breakpoints predicted as true somatic SVs in blue, whereas red bars represent the number of breakpoints predicted to as sequencing or mapping artefacts as a function of the confidence level.

**Supplementary Figure 5.**
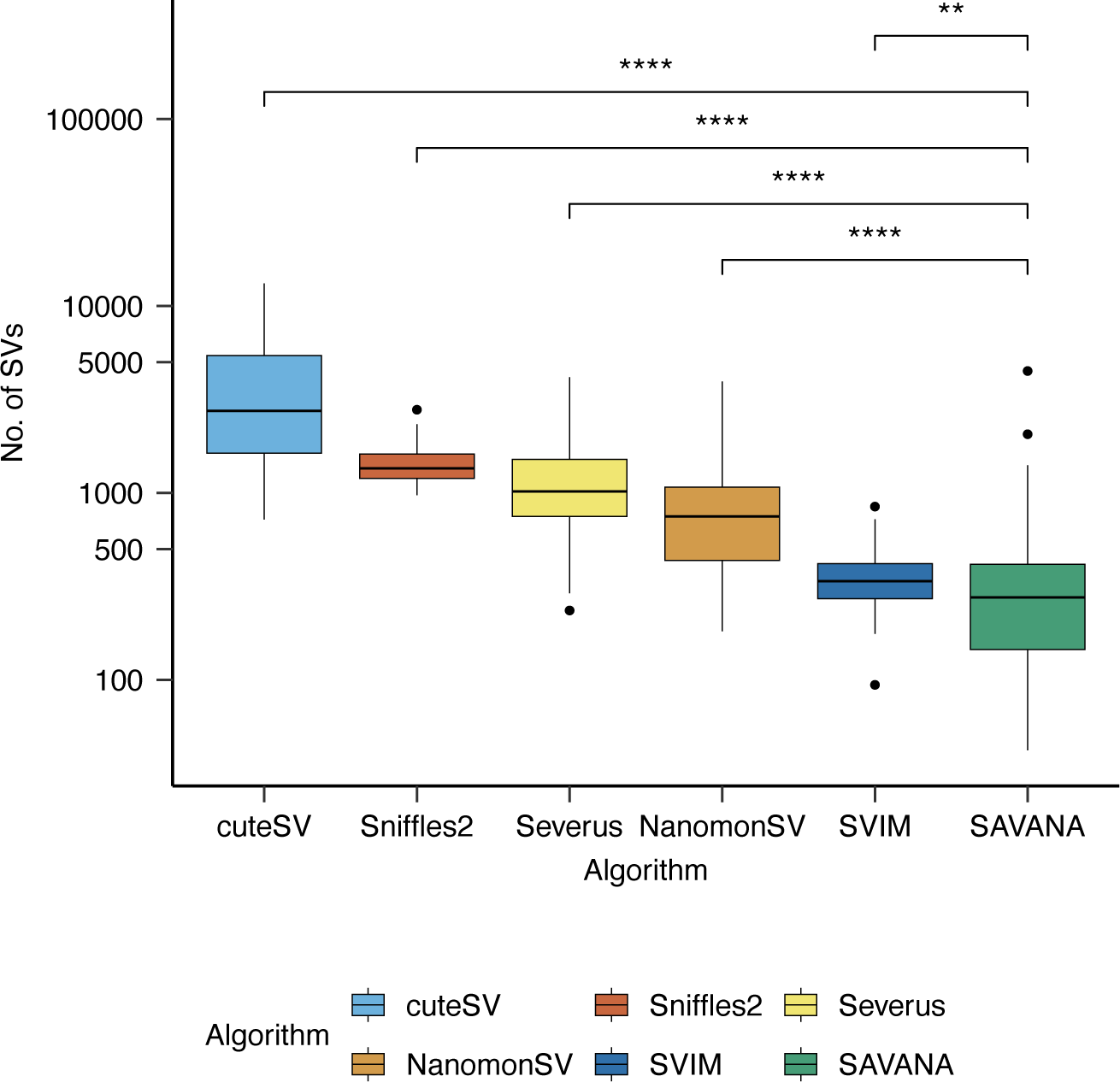
Total number of somatic SVs detected across the cohort using the SV detection algorithms benchmarked in this study. Significance was assessed using the two-sided Wilcoxon’s rank test; \*\**P* < 0.001, \*\*\*\**P* < 0.00001.

**Supplementary Figure 6.**
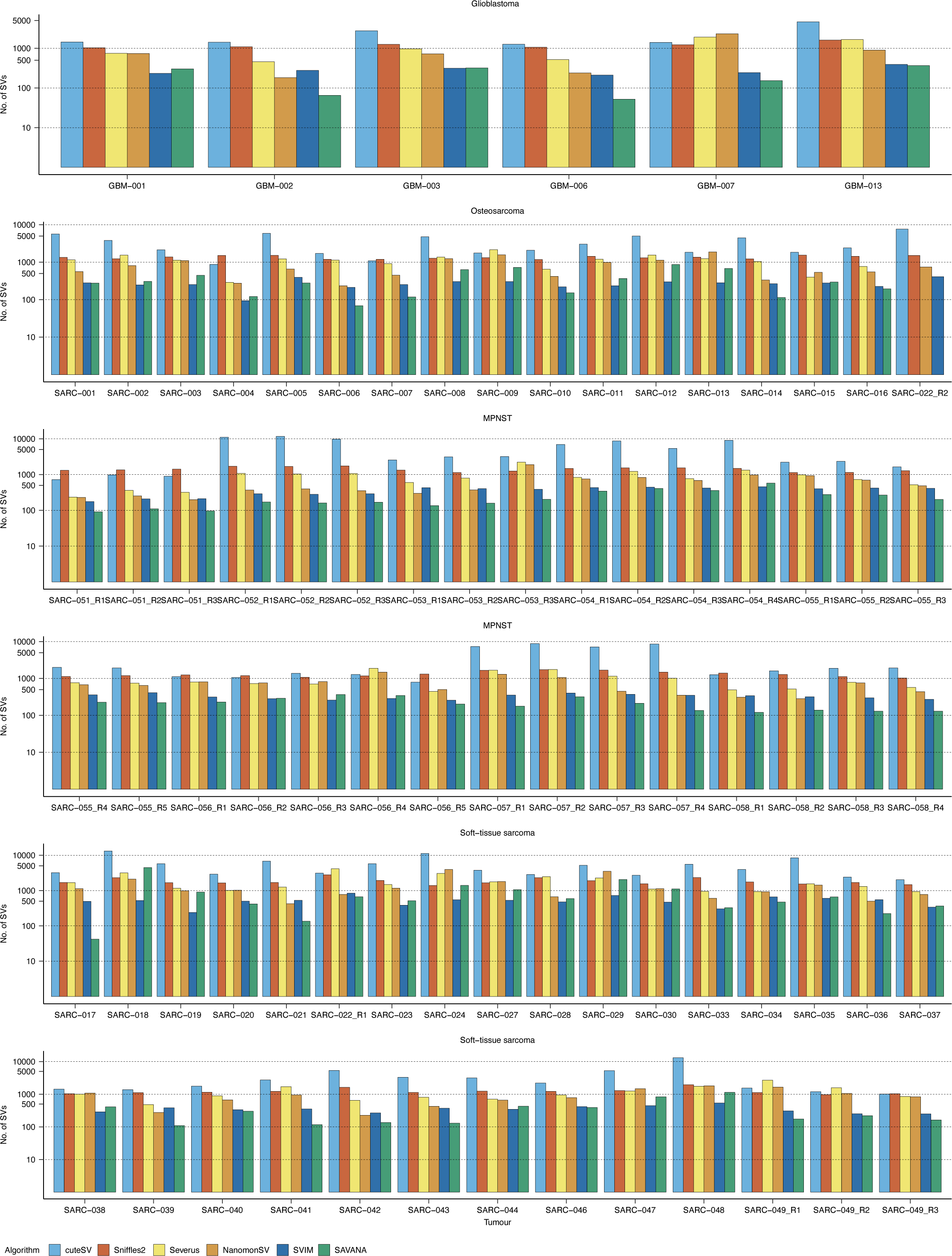
**Total number of somatic SVs detected in each sample by the algorithms benchmarked.**

**Supplementary Figure 7.**
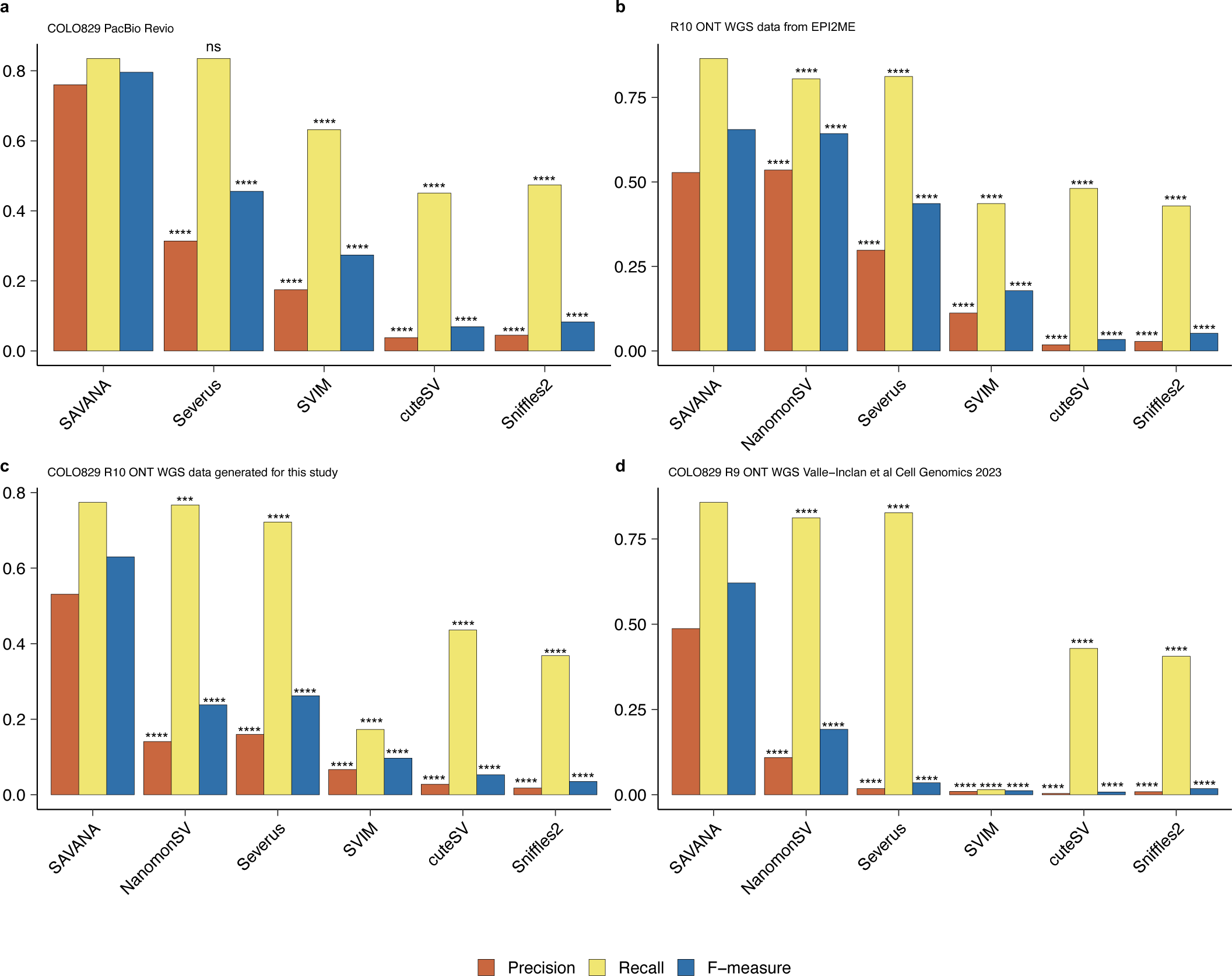
Benchmarking of SAVANA against existing SV detection algorithms using a truth set of 68 somatic SVs detected in the melanoma cell line COL0829 and its matched normal cell line COLO829BL. Benchmarking results using (**a**) WGS data generated using the Revio platform from PacBio; (**b**) nanopore WGS data generated by Oxford Nanopore Technologies using R10 PromethION flow cells; (**c**) nanopore WGS data generated for this study using R10 PromethION flow cells; and (**d**) nanopore WGS data generated by Valle-Inclán et al. Cell Genomics (2022) using R9 PromethION flow cells. Significance with respect to SAVANA was assessed using the two-sided Student’s *t*-test on 100 bootstrap resamples (\*\*\**P* < 0.0001; \*\*\*\**P* < 0.00001; ns: not significant).

**Supplementary Figure 8.**
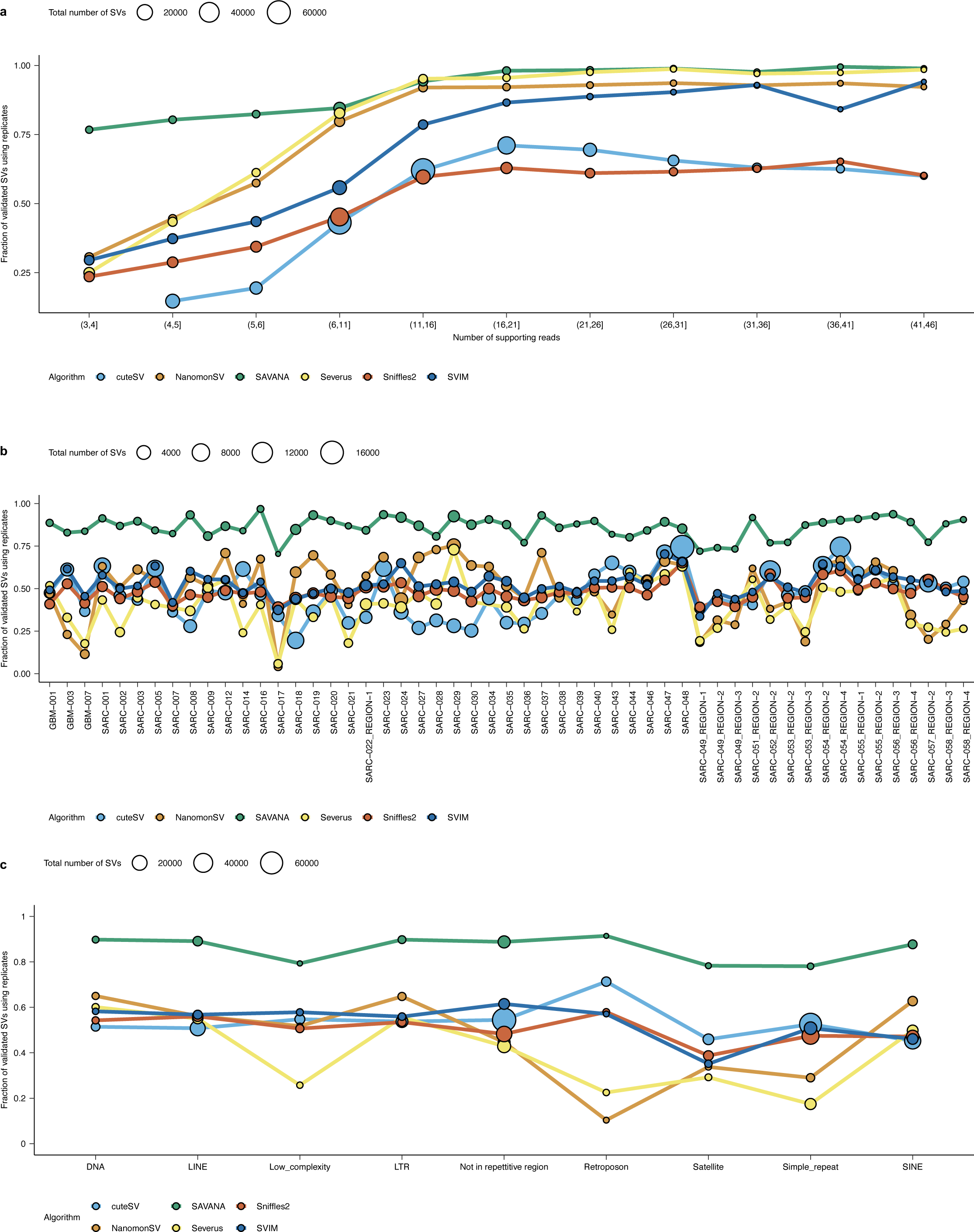
Total number of somatic SVs detected in each sample. (**a**) Fraction of somatic SVs detected in both replicates as a function of the number of reads supporting each SV. (**b**) Fraction of somatic SVs detected in both replicates for each sample. (**c**) Fraction of somatic SVs detected in both replicates across repetitive regions in the genome (GRCh38). The size of the dots in **a-c** represents the number of somatic SVs in each group. Panels **a-c** show the aggregated results for the 53 tumours with the highest sequencing depth.

**Supplementary Figure 9.**
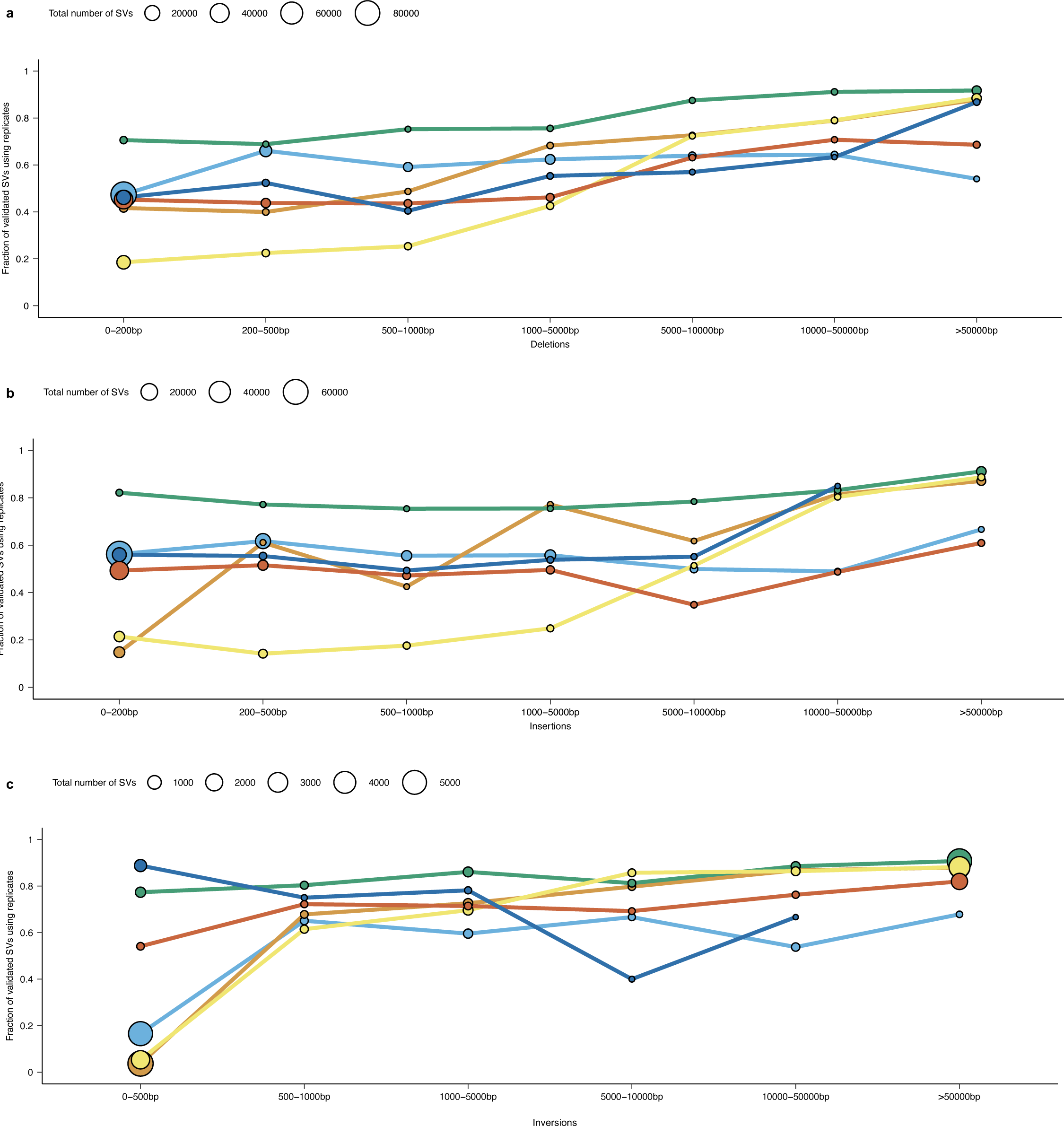
Fraction of replicated SVs as a function of SV size. Fraction of somatic deletions (**a**), insertions (**b**) and inversions (**c**) detected in both replicates as a function of SV size. The size of the dots in **a-c** represents the number of somatic SVs in each group. Panels **a-c** show the aggregated results for the 53 tumours with the highest sequencing depth.

**Supplementary Figure 10.**
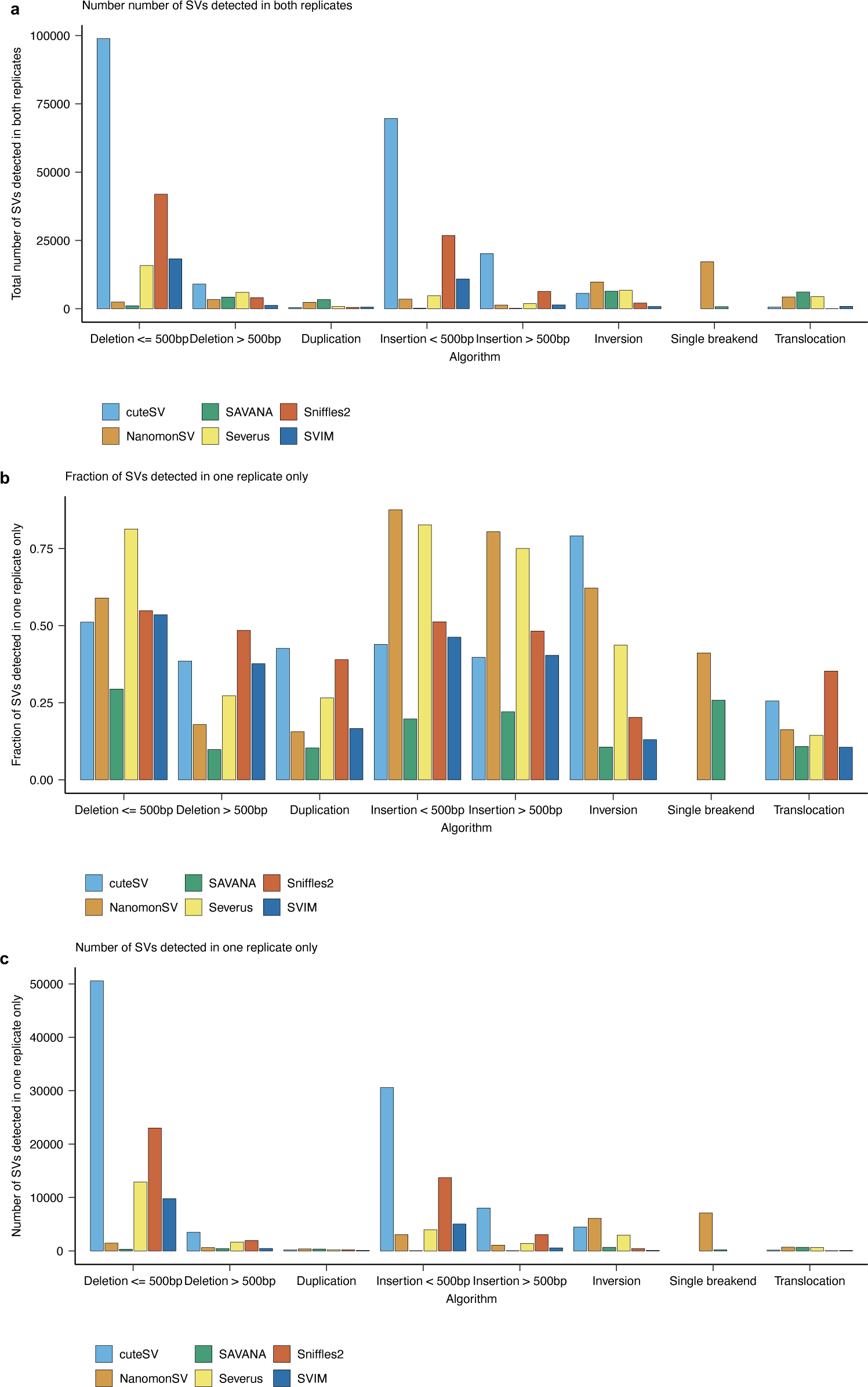
Analysis of somatic SVs detected across replicates stratified based on SV type. (**a**) Total number of SVs detected in all replicates. Fraction (**b**) and total number of SVs (**c**) detected in one replicate only. The bars in **a-c** show the aggregated data across all tumours analysed.

**Supplementary Figure 11.**
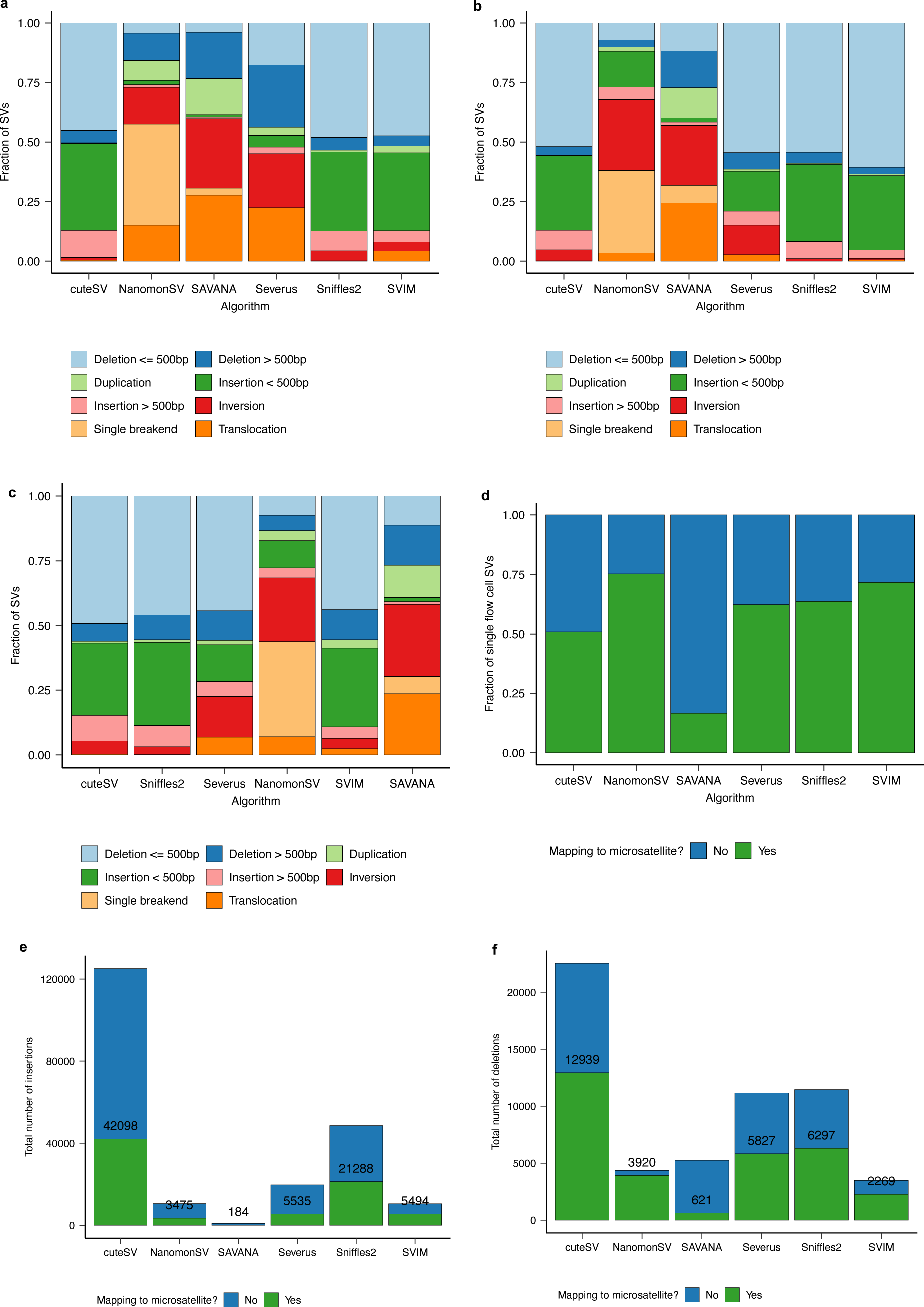
Analysis of the types of somatic SVs detected by the SV detection algorithms benchmarked. (**a**) Fraction of SVs detected in both replicates stratified based on the type of SV. (**b**) Fraction of SVs detected in one replicate only stratified based on the type of SV. (**c**) Distribution of SV types detected when the sequencing data from both flow cells was analysed jointly. (**d**) Fraction of SVs detected in one replicate only that mapped to microsatellite loci. Total number of insertions (**e**) and deletions (**f**) detected across all tumours when the sequencing data from all flow cells used to sequence each tumour sample were analysed jointly. The numbers on top of the bars in **e-f** indicate the total number of insertions and deletions, respectively, that mapped to microsatellite loci across all tumours analysed.

**Supplementary Figure 12.**
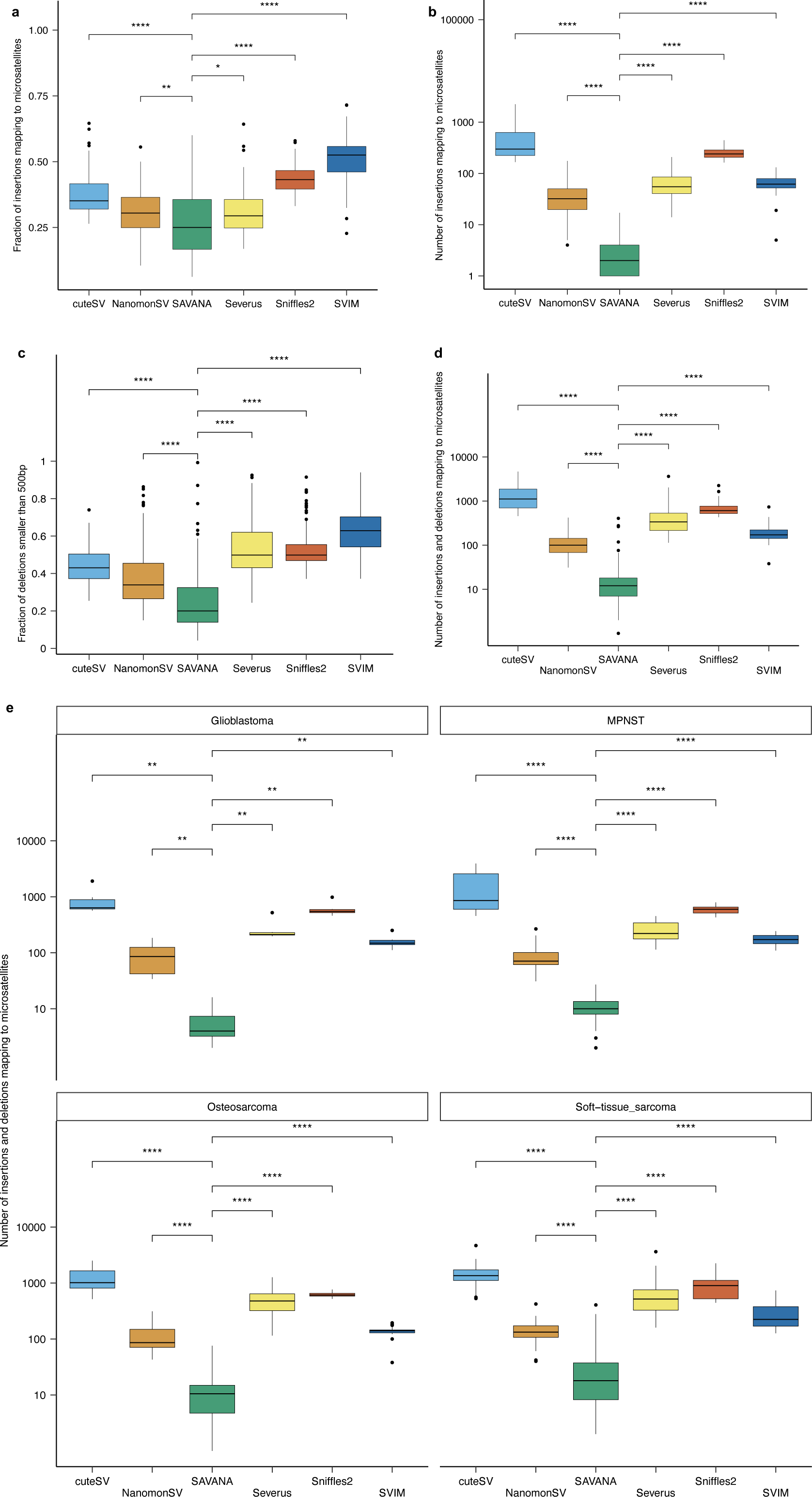
Analysis of the genomic distribution of deletions and insertions detected when analysing the tumour sequencing data from both flow cells jointly. Fraction (**a**) and total number (**b**) of insertions detected across all tumours analysed mapping to microsatellite loci. (**c**) Fraction of deletions smaller than 500bp detected across all tumours analysed mapping to microsatellite loci. (**d**) Fraction of deletions and insertions detected across all tumours analysed mapping to microsatellite loci. (**e**) Analysis of the fraction of deletions and insertions mapping to microsatellite loci stratified based on tumour type: glioblastomas, malignant peripheral nerve sheath tumours (MPNSTs), osteosarcomas, and soft-tissue sarcomas other than MPNSTs.

**Supplementary Figure 13.**
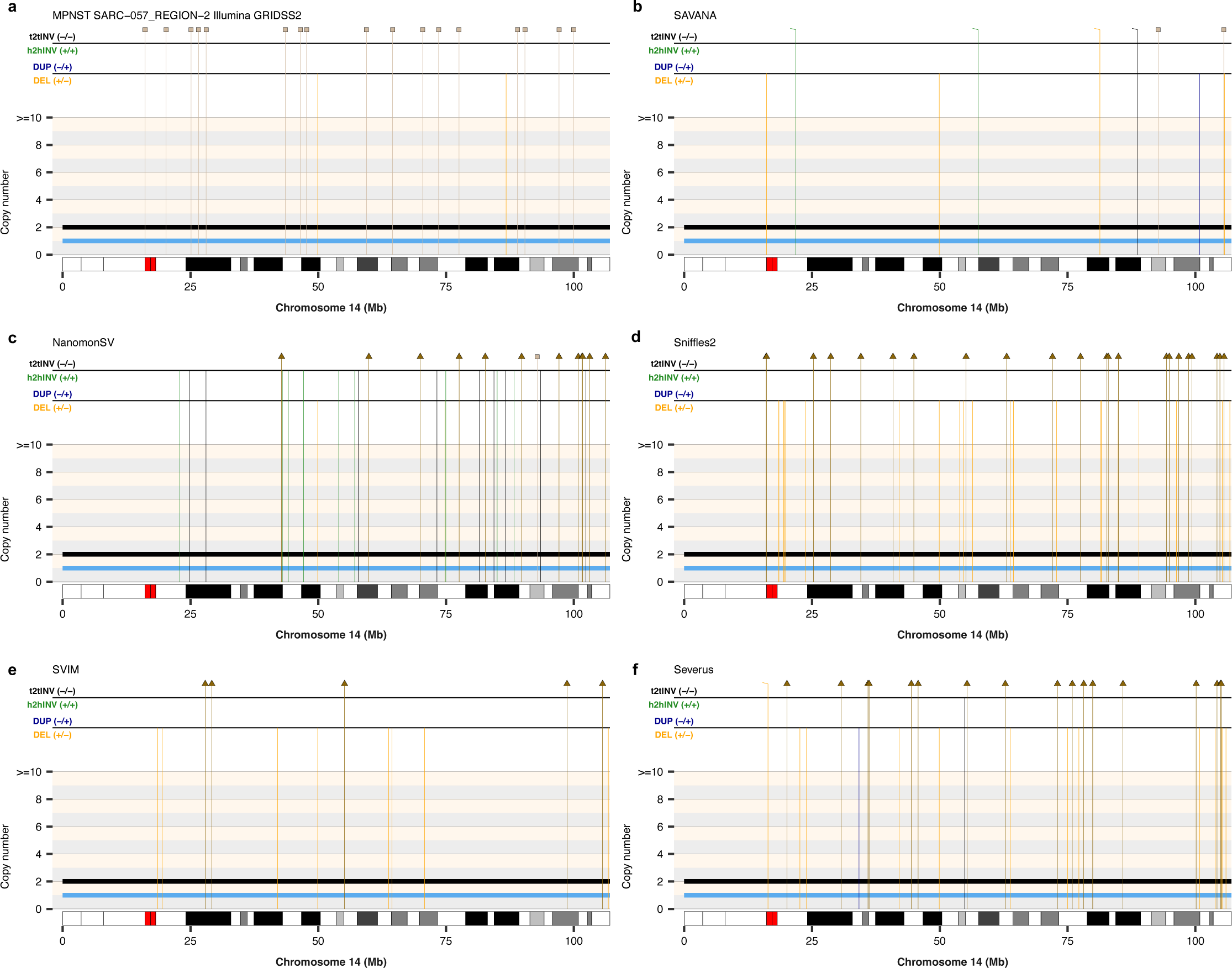
Comparison of the SVs detected in tumour region 2 from the MPNST SARC-057. (**a**) Somatic SVs and copy number profiles detected using GRIDSS2 and PURPLE in whole-genome short-read sequencing data. Somatic SVs detected in matched long-read nanopore whole-genome sequencing data using SAVANA (**b**), NanomonSV (**c**), Sniffles2 (**d**), SVIM (**e**) and Severus (**f**). The copy number profiles shown in **a-f** were calculated using PURPLE and the short-read sequencing data. The total and minor allele copy-number data in **a-f** are represented in black and blue, respectively. DEL, deletion-like rearrangement; DUP, duplication-like rearrangement; h2hINV, head-to-head inversion; t2tINV, tail-to-tail inversion. Lines with a square at the top represent single breakends, and lines with arrowheads mark insertions.

**Supplementary Figure 14.**
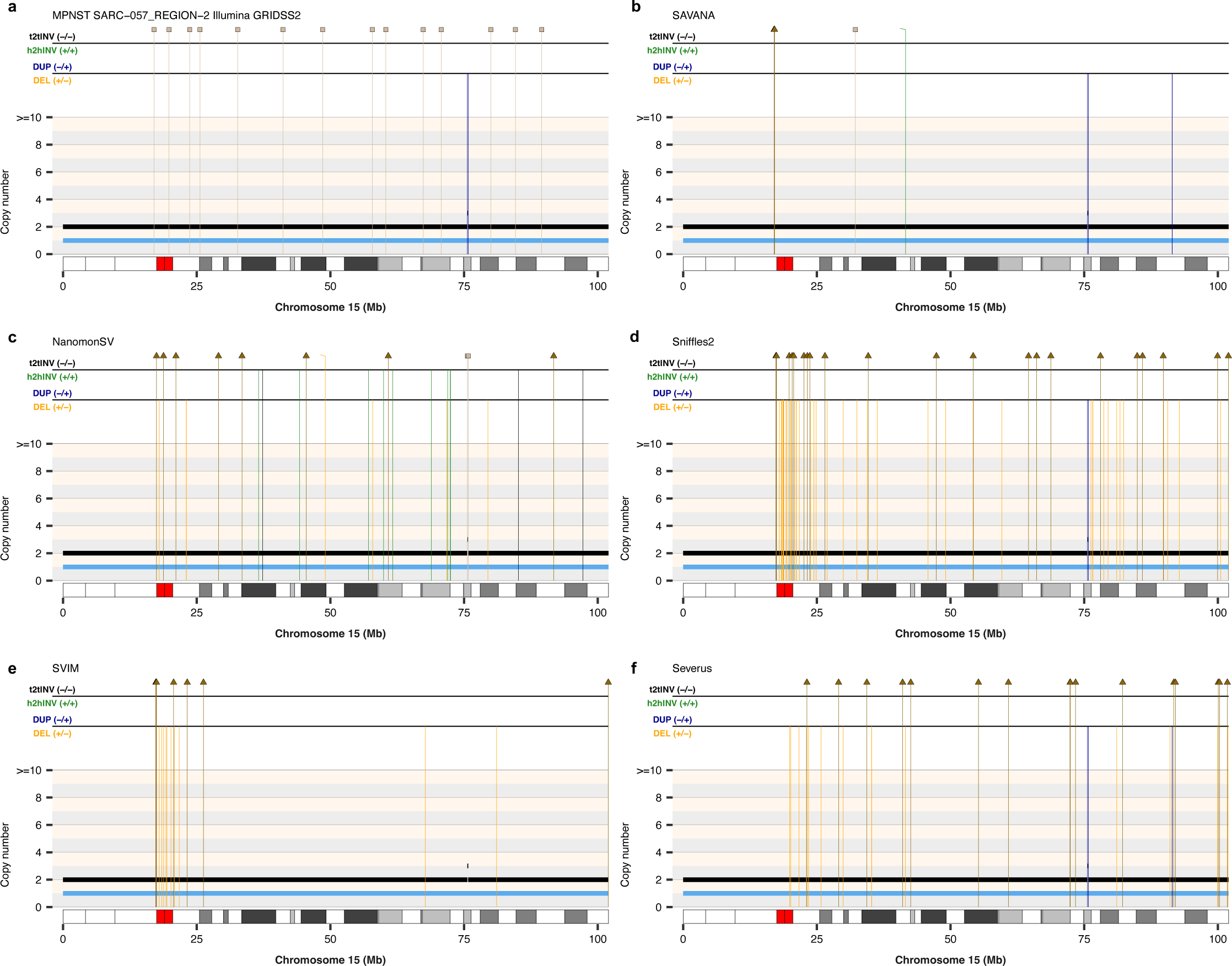
Comparison of the SVs detected in tumour region 2 from the MPNST SARC-057. (**a**) Somatic SVs and copy number profiles detected using GRIDSS2 and PURPLE in whole-genome short-read sequencing data. Somatic SVs detected in matched long-read nanopore whole-genome sequencing data using SAVANA (**b**), NanomonSV (**c**), Sniffles2 (**d**), SVIM (**e**) and Severus (**f**). The copy number profiles shown in **a-f** were calculated using PURPLE and the short-read sequencing data. The total and minor allele copy-number data in **a-f** are represented in black and blue, respectively. DEL, deletion-like rearrangement; DUP, duplication-like rearrangement; h2hINV, head-to-head inversion; t2tINV, tail-to-tail inversion. Lines with a square at the top represent single breakends, and lines with arrowheads mark insertions.

**Supplementary Figure 15.**
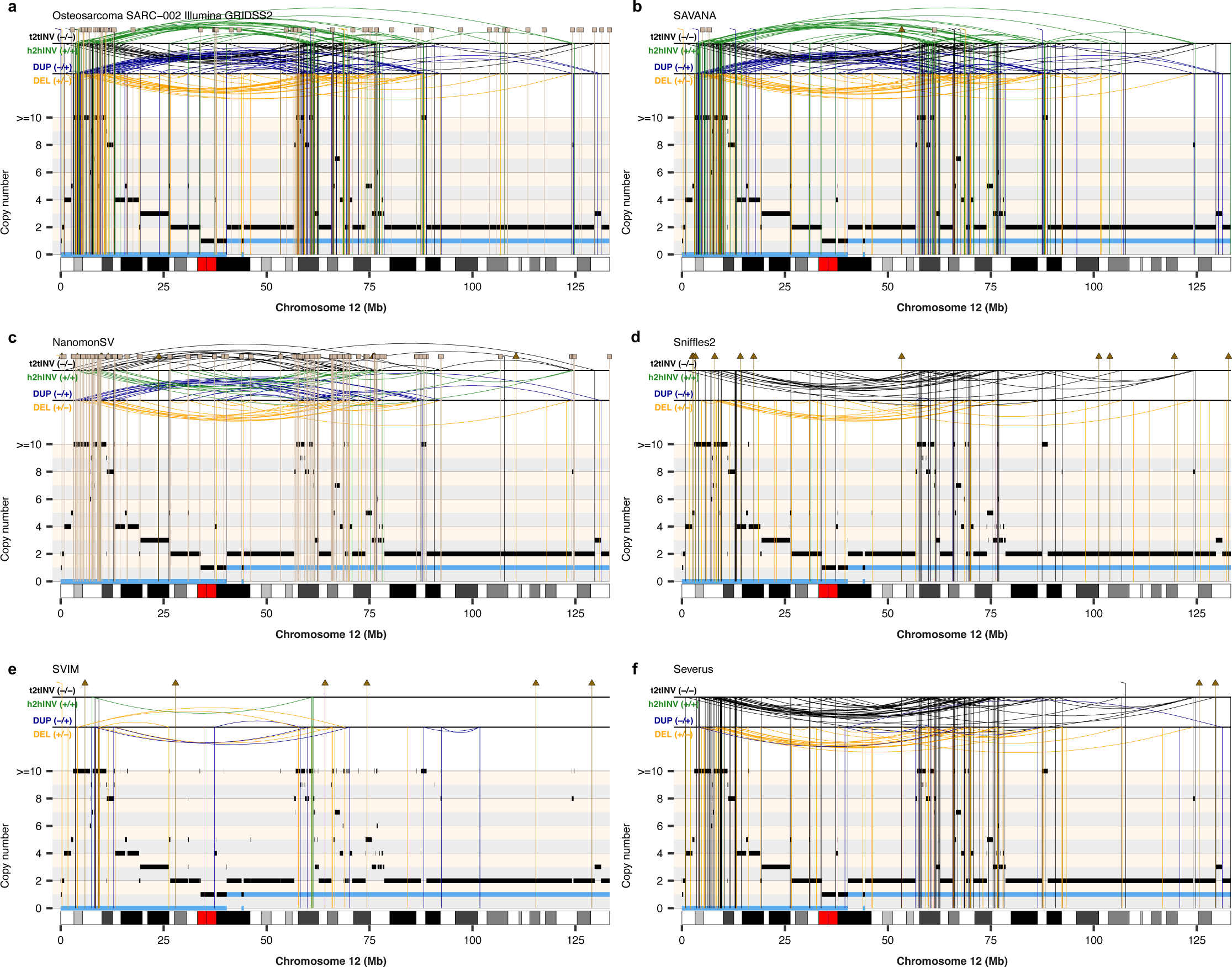
Comparison of the SVs detected in the osteosarcoma SARC-002. (**a**) Somatic SVs and copy number profiles detected using GRIDSS2 and PURPLE in whole-genome short-read sequencing data. Somatic SVs detected in matched long-read nanopore whole-genome sequencing data using SAVANA (**b**), NanomonSV (**c**), Sniffles2 (**d**), SVIM (**e**) and Severus (**f**). The copy number profiles shown in **a-f** were calculated using PURPLE and the short-read sequencing data. The total and minor allele copy-number data in **a-f** are represented in black and blue, respectively. DEL, deletion-like rearrangement; DUP, duplication-like rearrangement; h2hINV, head-to-head inversion; t2tINV, tail-to-tail inversion. Lines with a square at the top represent single breakends, and lines with arrowheads mark insertions.

**Supplementary Figure 16.**
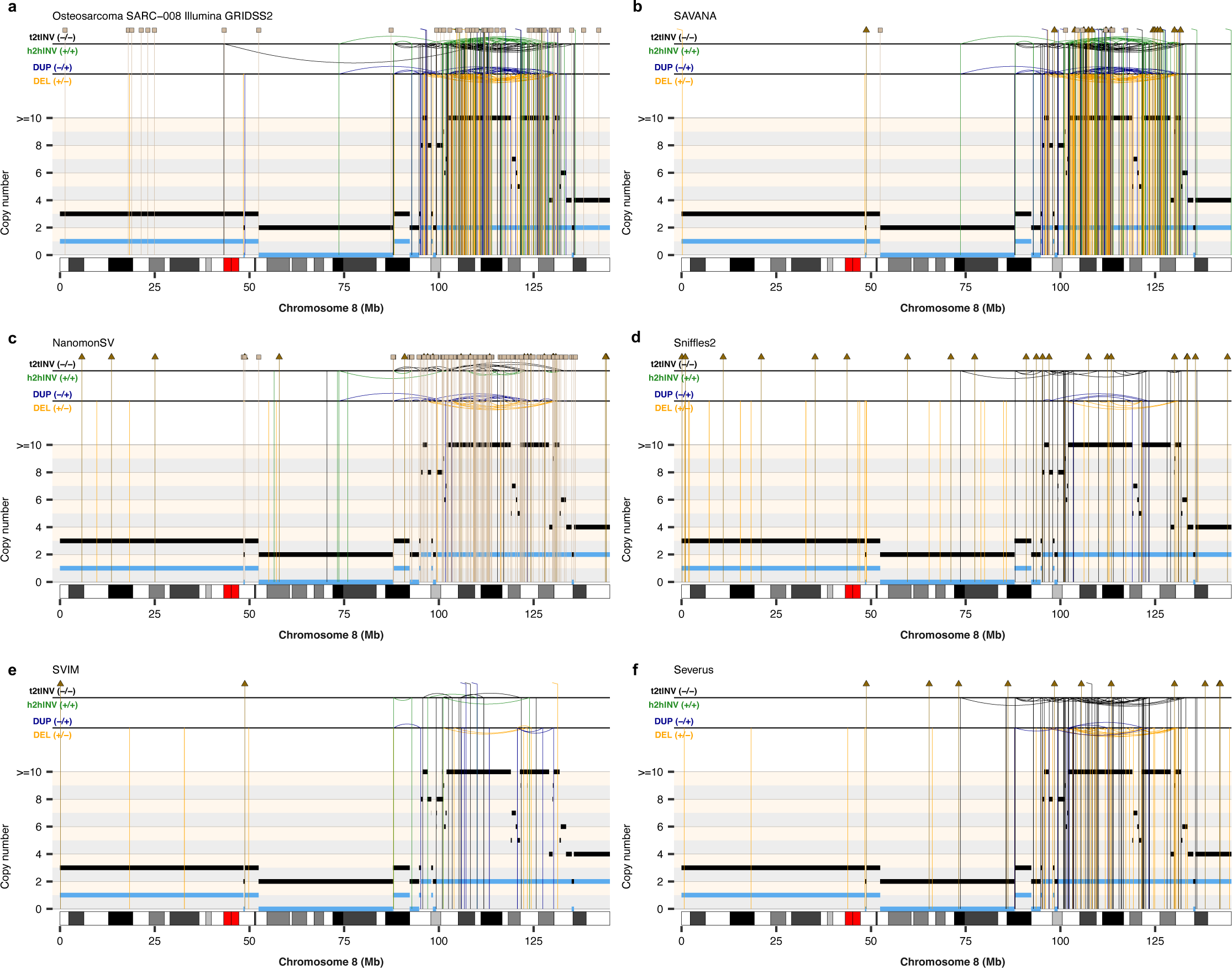
Comparison of the SVs detected in the osteosarcoma SARC-008. (**a**) Somatic SVs and copy number profiles detected using GRIDSS2 and PURPLE in whole-genome short-read sequencing data. Somatic SVs detected in matched long-read nanopore whole-genome sequencing data using SAVANA (**b**), NanomonSV (**c**), Sniffles2 (**d**), SVIM (**e**) and Severus (**f**). The copy number profiles shown in **a-f** were calculated using PURPLE and the short-read sequencing data. The total and minor allele copy-number data in **a-f** are represented in black and blue, respectively. DEL, deletion-like rearrangement; DUP, duplication-like rearrangement; h2hINV, head-to-head inversion; t2tINV, tail-to-tail inversion. Lines with a square at the top represent single breakends, and lines with arrowheads mark insertions.

**Supplementary Figure 17.**
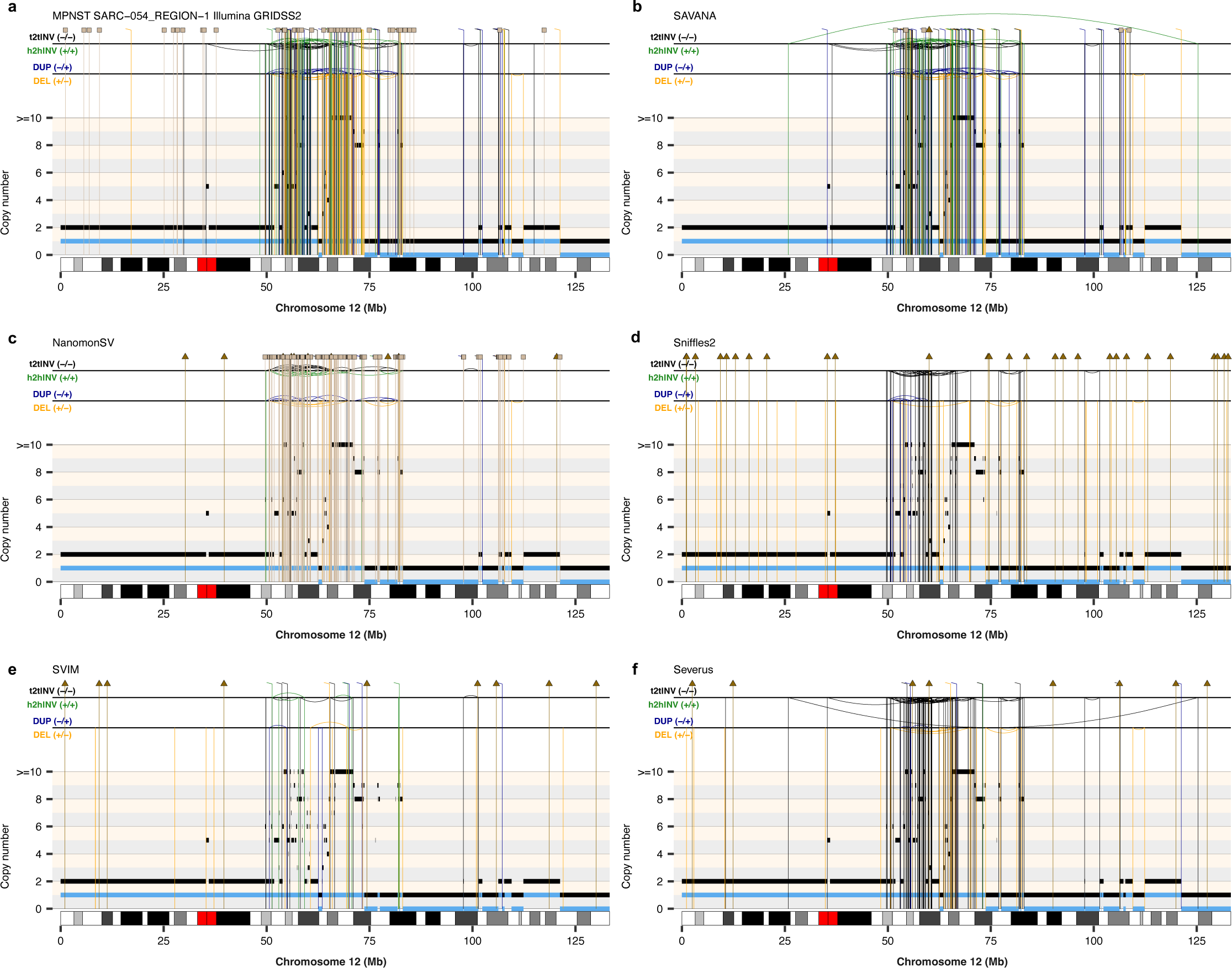
Comparison of the SVs detected in the myxofibrosarcoma SARC-054. (**a**) Somatic SVs and copy number profiles detected using GRIDSS2 and PURPLE in whole-genome short-read sequencing data. Somatic SVs detected in matched long-read nanopore whole-genome sequencing data using SAVANA (**b**), NanomonSV (**c**), Sniffles2 (**d**), SVIM (**e**) and Severus (**f**). The copy number profiles shown in **a-f** were calculated using PURPLE and the short-read sequencing data. The total and minor allele copy-number data in **a-f** are represented in black and blue, respectively. DEL, deletion-like rearrangement; DUP, duplication-like rearrangement; h2hINV, head-to-head inversion; t2tINV, tail-to-tail inversion. Lines with a square at the top represent single breakends, and lines with arrowheads mark insertions.

**Supplementary Figure 18.**
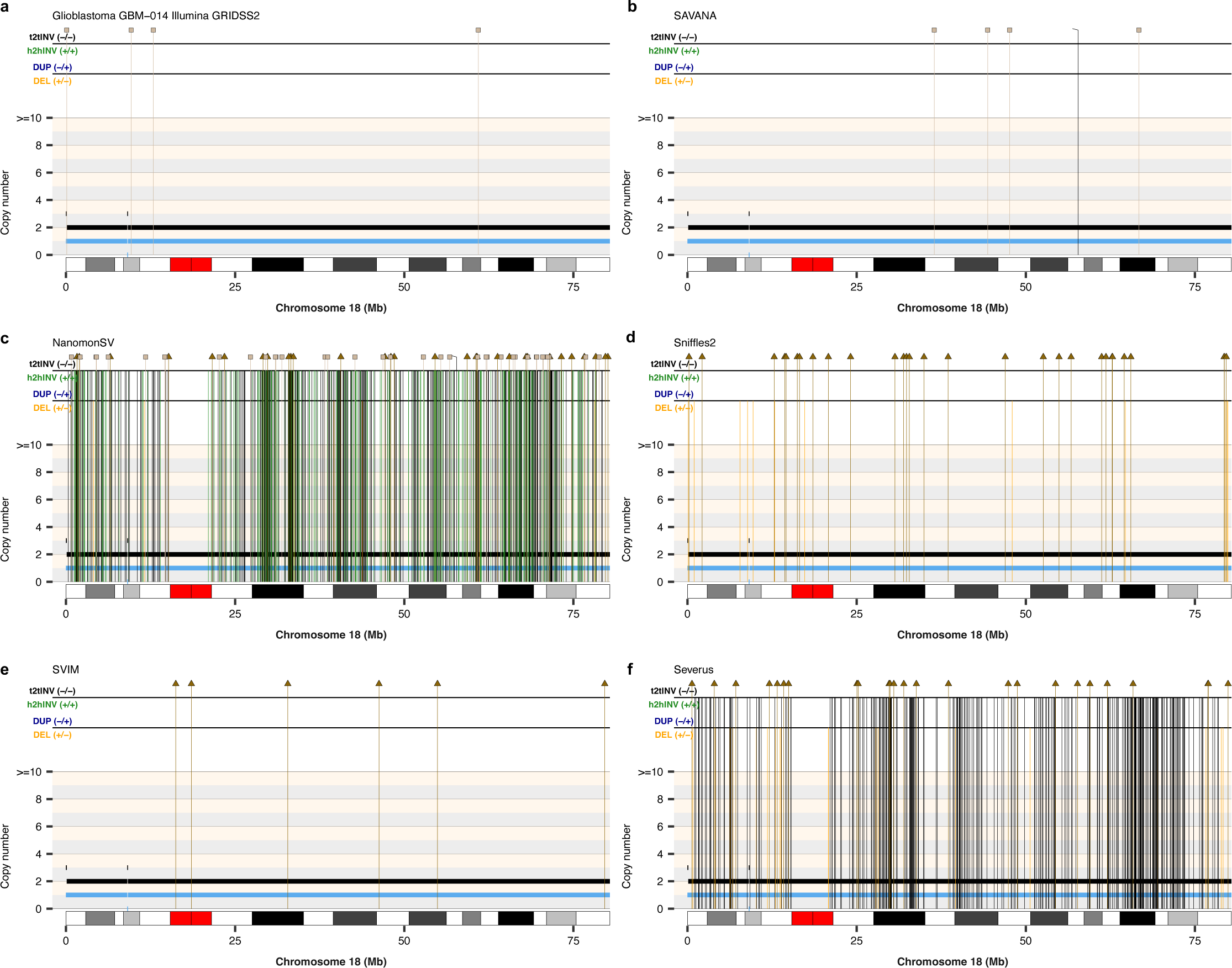
Comparison of the SVs detected in the glioblastoma GBM-014. (**a**) Somatic SVs and copy number profiles detected using GRIDSS2 and PURPLE in whole-genome short-read sequencing data. Somatic SVs detected in matched long-read nanopore whole-genome sequencing data using SAVANA (**b**), NanomonSV (**c**), Sniffles2 (**d**), SVIM (**e**) and Severus (**f**). The copy number profiles shown in **a-f** were calculated using PURPLE and the short-read sequencing data. The total and minor allele copy-number data in **a-f** are represented in black and blue, respectively. DEL, deletion-like rearrangement; DUP, duplication-like rearrangement; h2hINV, head-to-head inversion; t2tINV, tail-to-tail inversion. Lines with a square at the top represent single breakends, and lines with arrowheads mark insertions.

**Supplementary Figure 19.**
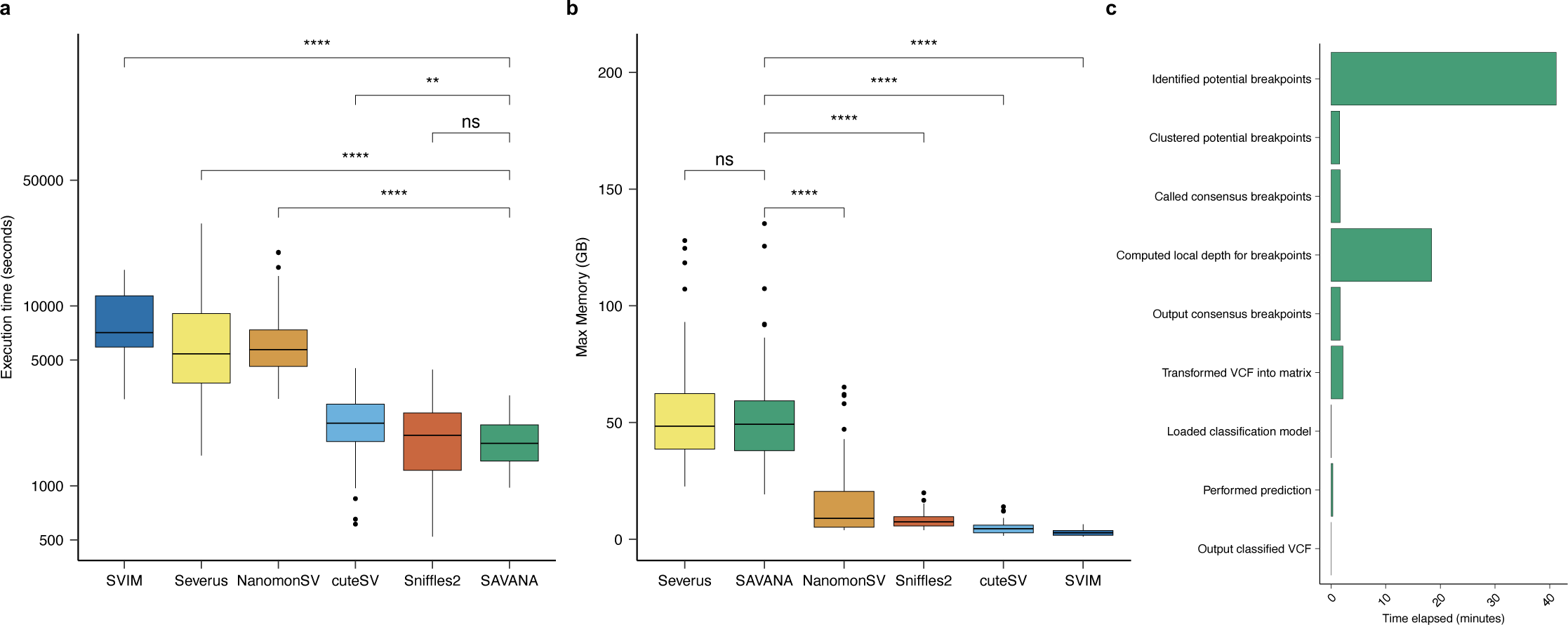
Comparison of the execution time (a) and memory use (b) between SAVANA and existing SV detection algorithms across the cohort. (**c**) Running time for each step of SAVANA. Significance in **a-b** was assessed using the two-sided Wilcoxon’s rank test (\*\**P* < 0.001; \*\*\*\**P* < 0.00001).

**Supplementary Figure 20.**
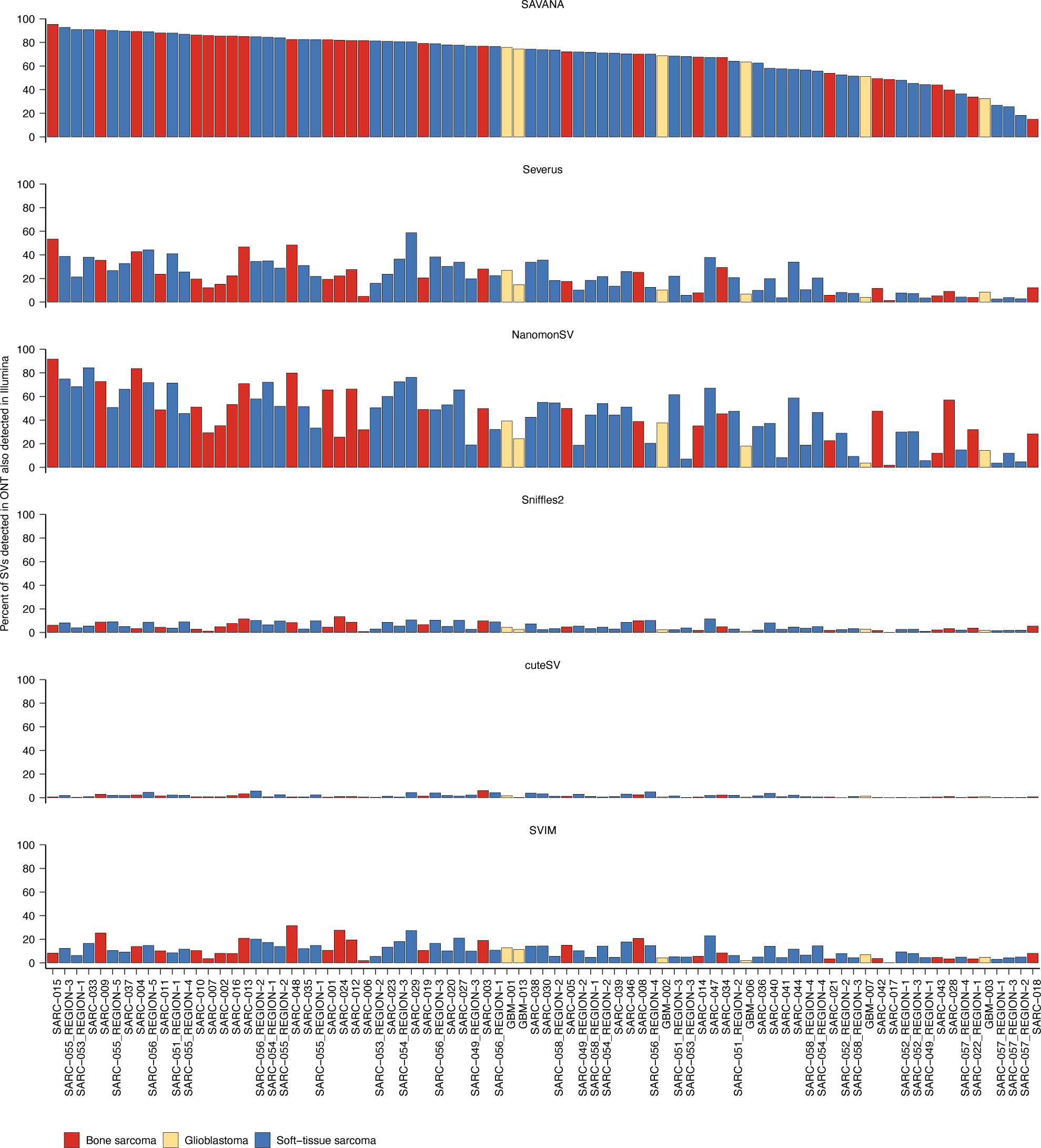
Comparison between short and long read sequencing data for somatic SV detection. Each bar reports the percent of somatic SVs detected in nanopore long-read WGS data that are also detected in matched Illumina WGS data.

**Supplementary Figure 21.**
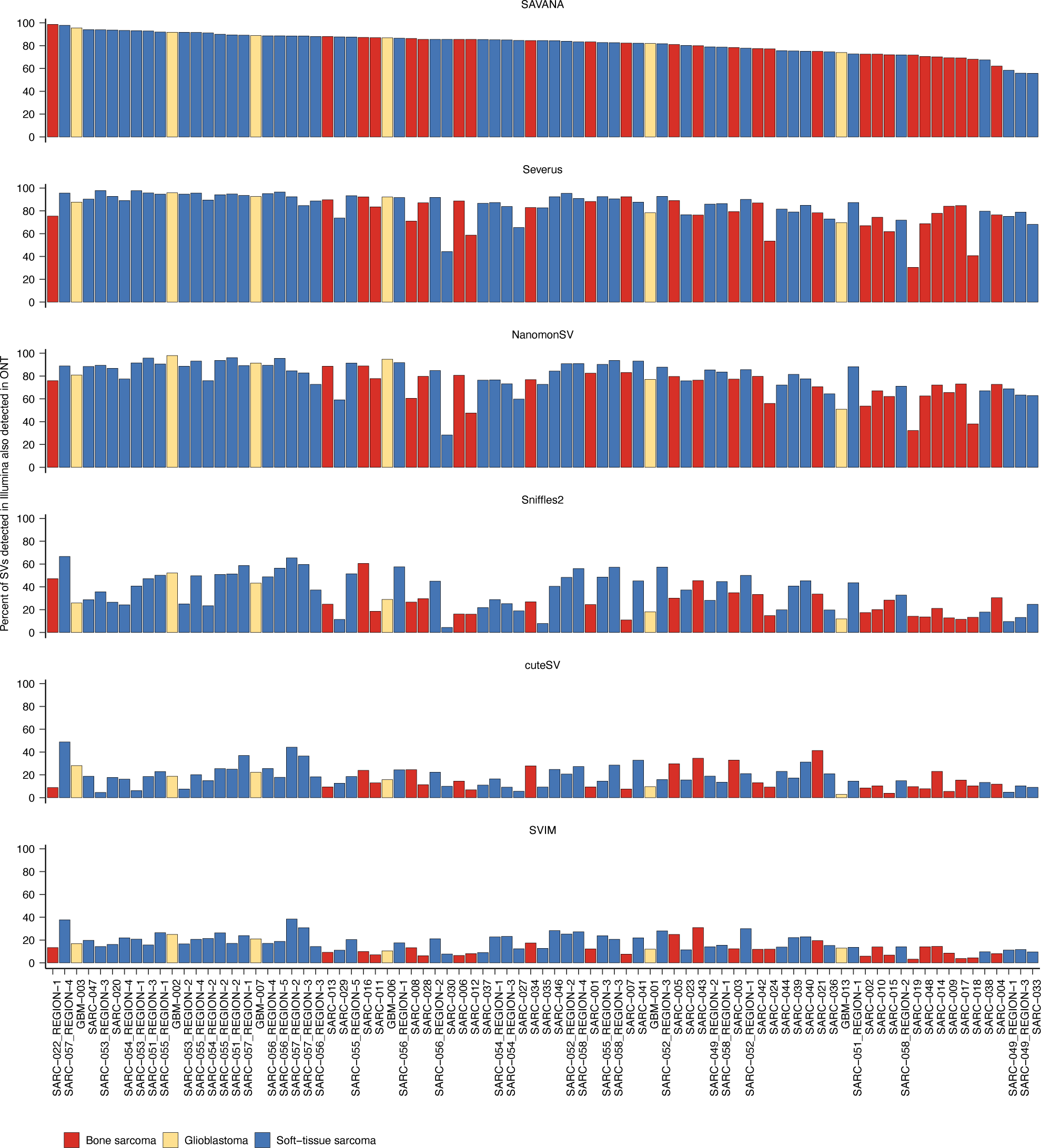
Comparison between short and long read sequencing data for somatic SV detection. Each bar reports the percent of somatic SVs detected in Illumina WGS data that are also detected in matched nanopore long-read WGS data.

**Supplementary Figure 22.**
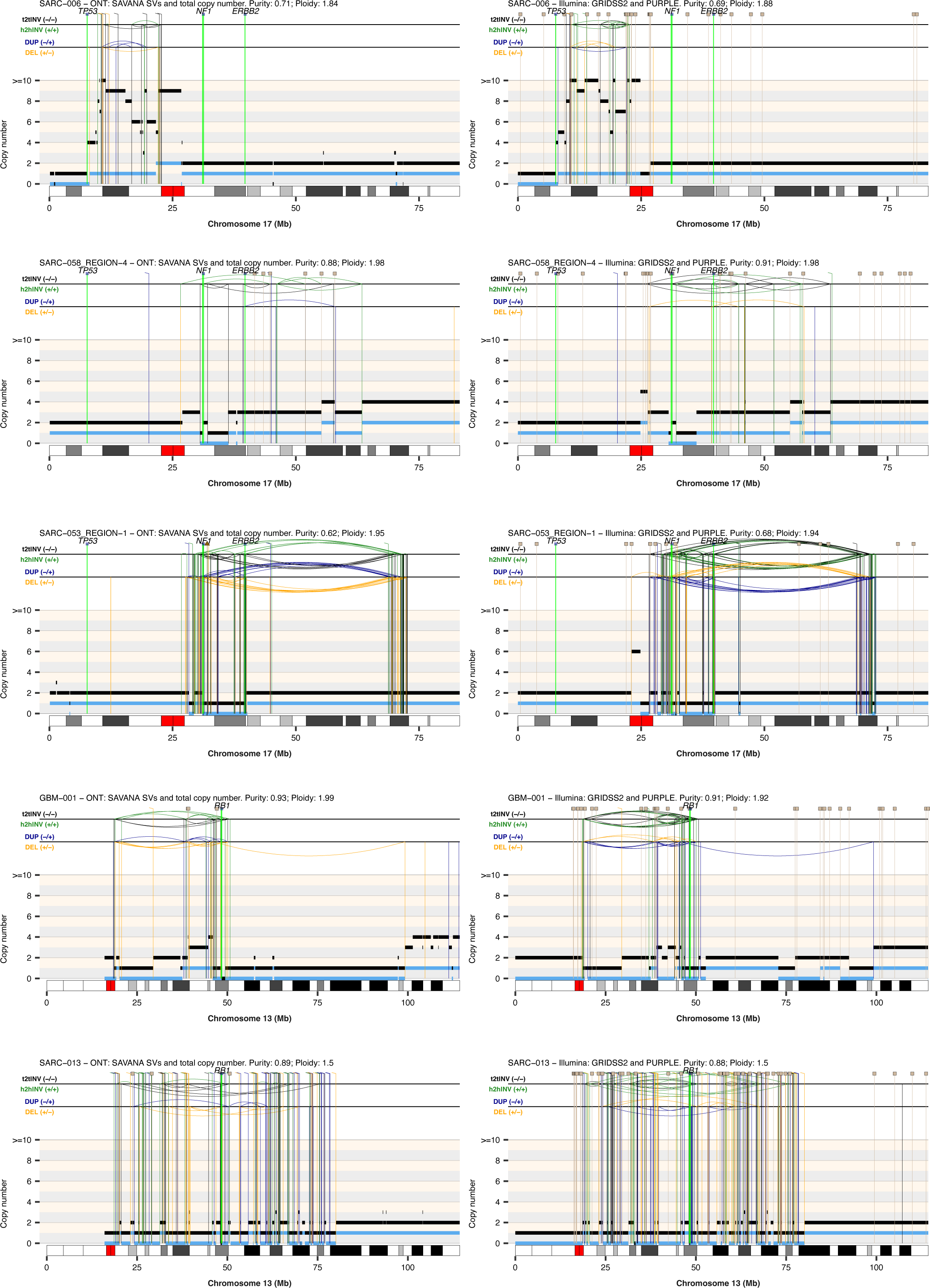
Somatic rearrangement and copy number profiles calculated using short-read data analyzed with GRIDSS and PURPLE against long-read data analyzed using SAVANA. (**Left**) Somatic SVs and SCNAs detected in matched long-read nanopore whole-genome sequencing data using SAVANA. (**Right**) Somatic SVs and copy number profiles detected using GRIDSS2 and PURPLE in whole-genome short-read sequencing data. The total and minor allele copy-number data are represented in black and blue, respectively. DEL, deletion-like rearrangement; DUP, duplication-like rearrangement; h2hINV, head-to-head inversion; t2tINV, tail-to-tail inversion. Lines with a square at the top represent single breakends, and lines with arrowheads mark insertions.

**Supplementary Figure 23.**
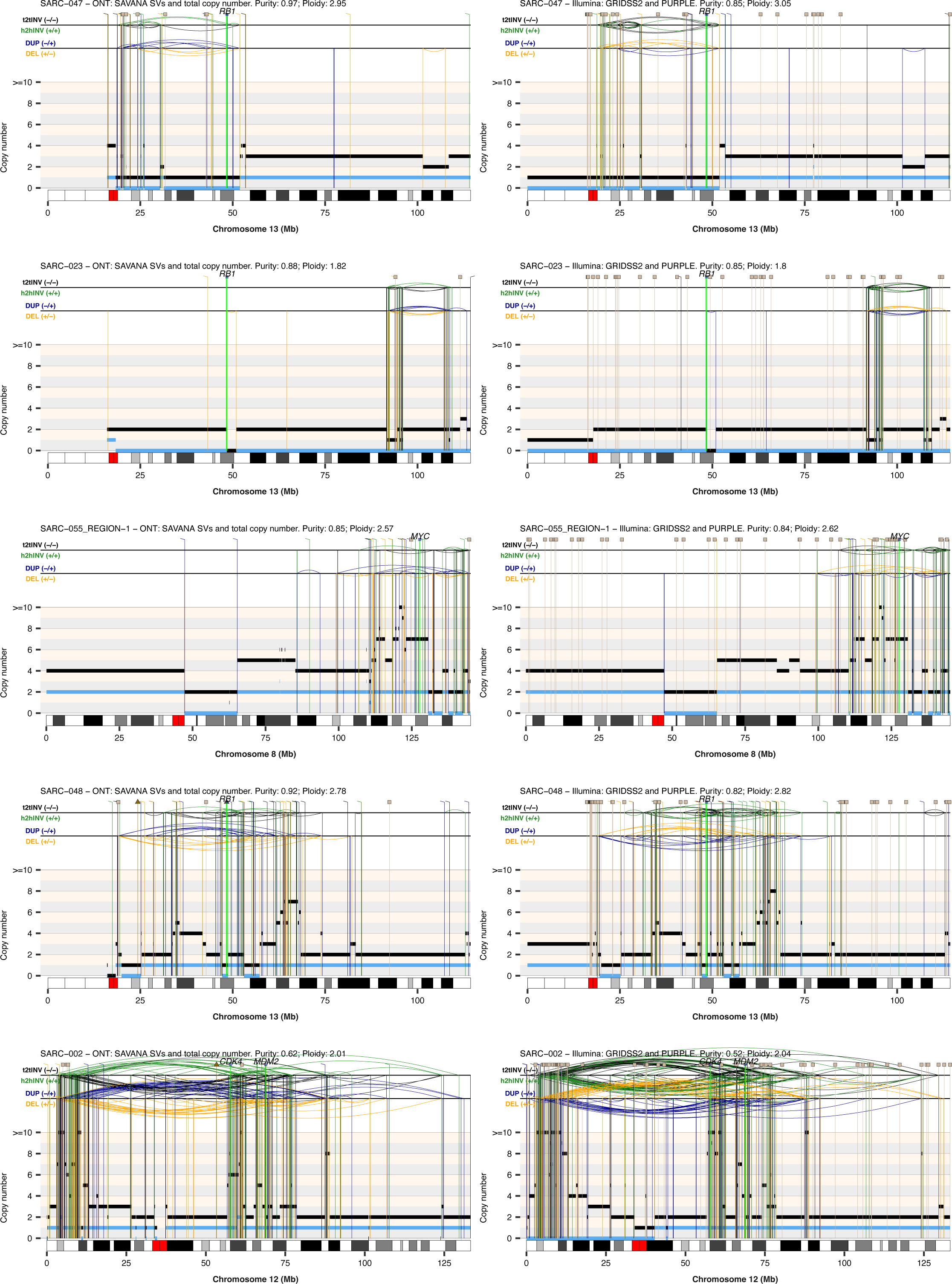
Somatic rearrangement and copy number profiles calculated using short-read data analyzed with GRIDSS and PURPLE against long-read data analyzed using SAVANA. (**Left**) Somatic SVs and SCNAs detected in matched long-read nanopore whole-genome sequencing data using SAVANA. (**Right**) Somatic SVs and copy number profiles detected using GRIDSS2 and PURPLE in whole-genome short-read sequencing data. The total and minor allele copy-number data are represented in black and blue, respectively. DEL, deletion-like rearrangement; DUP, duplication-like rearrangement; h2hINV, head-to-head inversion; t2tINV, tail-to-tail inversion. Lines with a square at the top represent single breakends, and lines with arrowheads mark insertions.

**Supplementary Figure 24.**
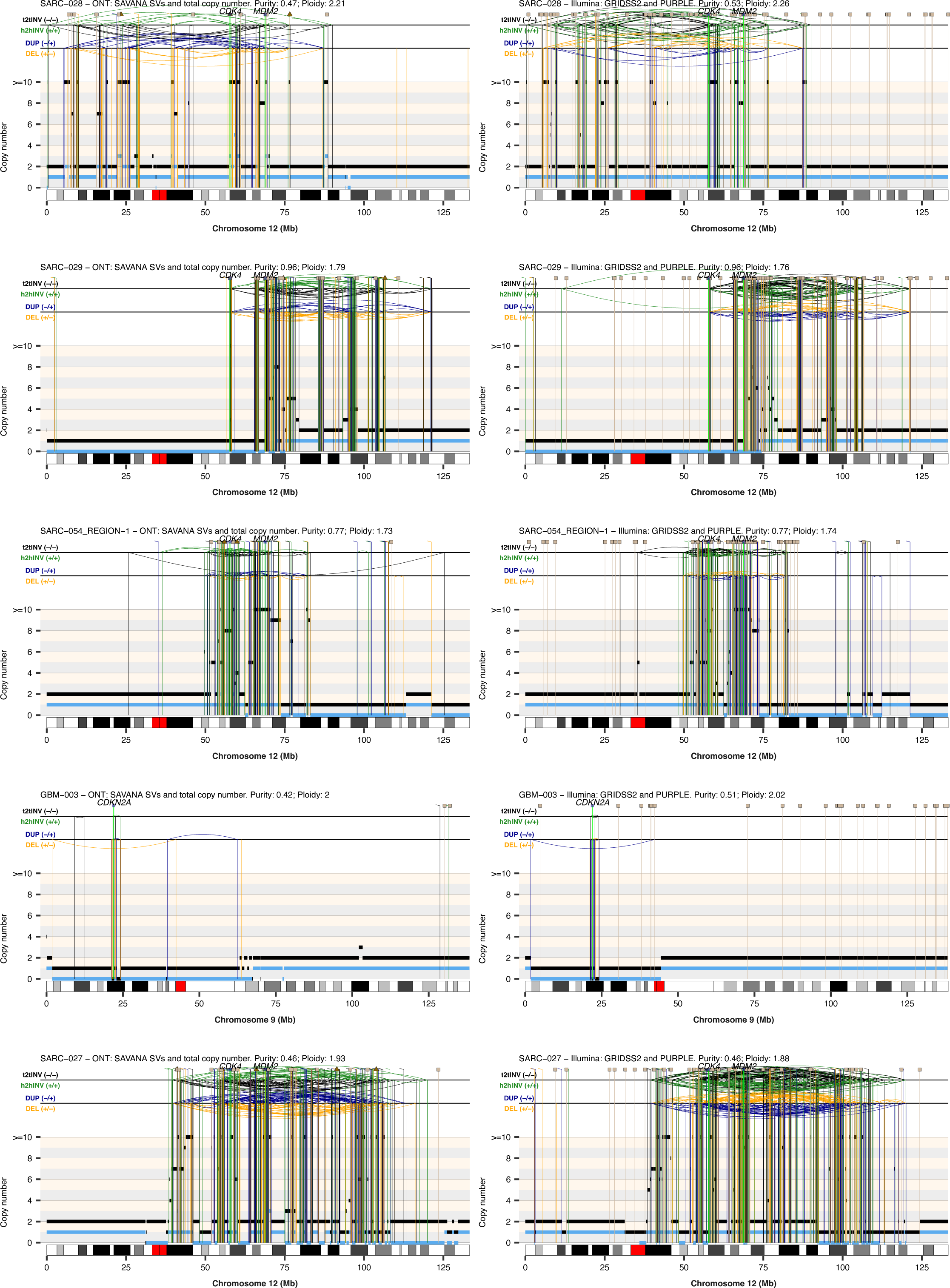
Somatic rearrangement and copy number profiles calculated using short-read data analyzed with GRIDSS and PURPLE against long-read data analyzed using SAVANA. (**Left**) Somatic SVs and SCNAs detected in matched long-read nanopore whole-genome sequencing data using SAVANA. (**Right**) Somatic SVs and copy number profiles detected using GRIDSS2 and PURPLE in whole-genome short-read sequencing data. The total and minor allele copy-number data are represented in black and blue, respectively. DEL, deletion-like rearrangement; DUP, duplication-like rearrangement; h2hINV, head-to-head inversion; t2tINV, tail-to-tail inversion. Lines with a square at the top represent single breakends, and lines with arrowheads mark insertions.

**Supplementary Figure 25.**
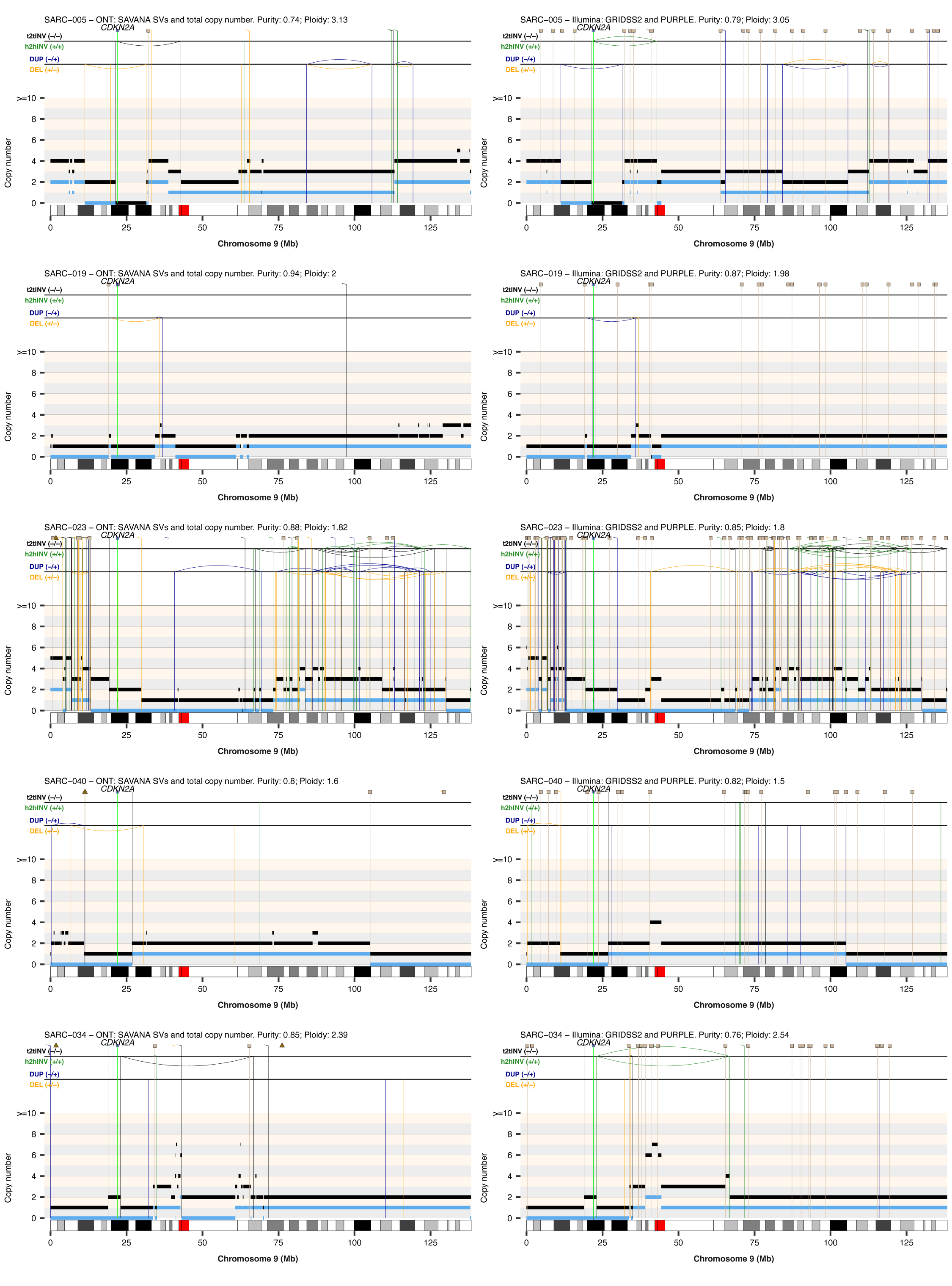
Somatic rearrangement and copy number profiles calculated using short-read data analyzed with GRIDSS and PURPLE against long-read data analyzed using SAVANA. (**Left**) Somatic SVs and SCNAs detected in matched long-read nanopore whole-genome sequencing data using SAVANA. (**Right**) Somatic SVs and copy number profiles detected using GRIDSS2 and PURPLE in whole-genome short-read sequencing data. The total and minor allele copy-number data are represented in black and blue, respectively. DEL, deletion-like rearrangement; DUP, duplication-like rearrangement; h2hINV, head-to-head inversion; t2tINV, tail-to-tail inversion. Lines with a square at the top represent single breakends, and lines with arrowheads mark insertions.

**Supplementary Figure 26.**
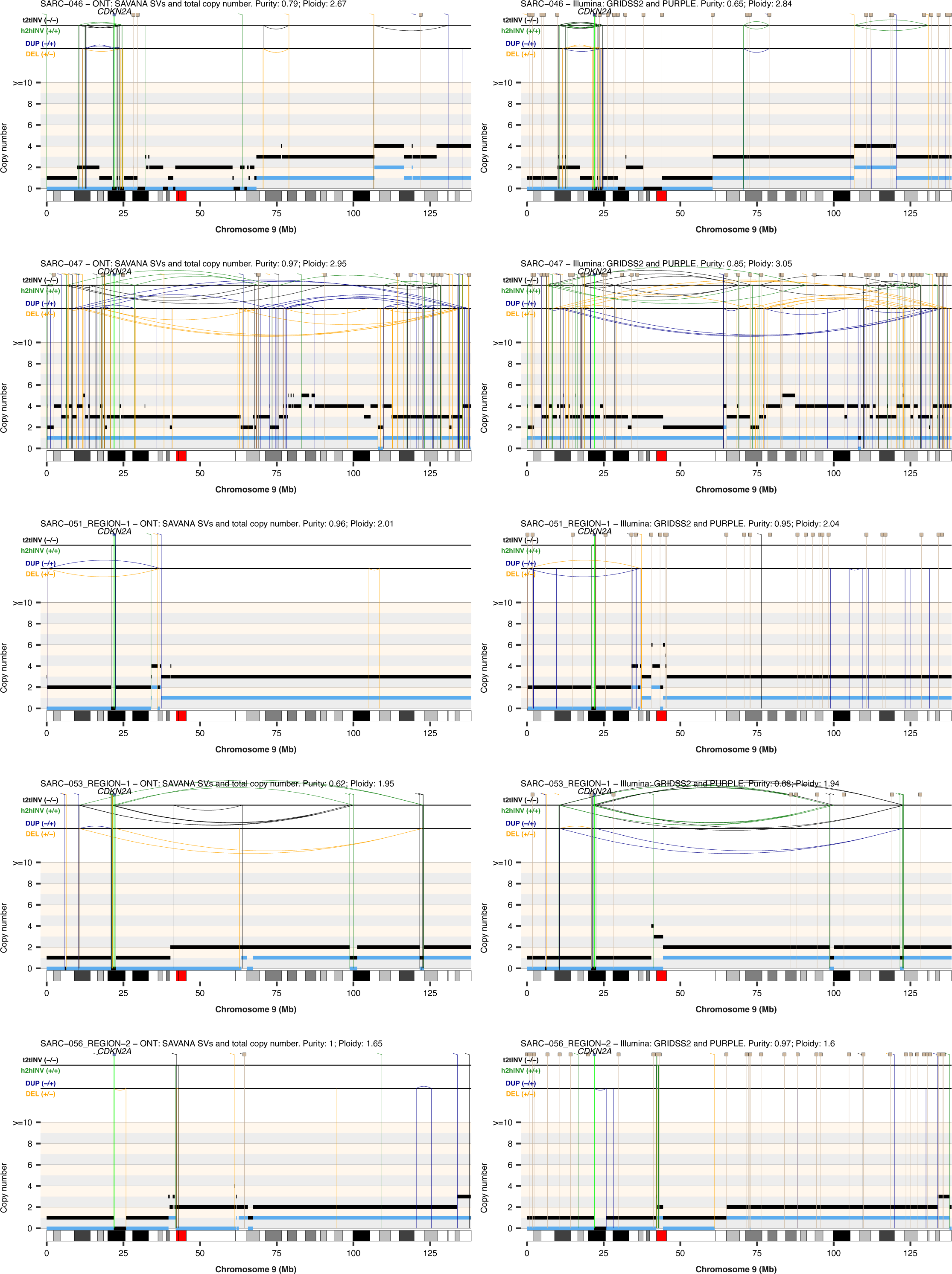
Somatic rearrangement and copy number profiles calculated using short-read data analyzed with GRIDSS and PURPLE against long-read data analyzed using SAVANA. (**Left**) Somatic SVs and SCNAs detected in matched long-read nanopore whole-genome sequencing data using SAVANA. (**Right**) Somatic SVs and copy number profiles detected using GRIDSS2 and PURPLE in whole-genome short-read sequencing data. The total and minor allele copy-number data are represented in black and blue, respectively. DEL, deletion-like rearrangement; DUP, duplication-like rearrangement; h2hINV, head-to-head inversion; t2tINV, tail-to-tail inversion. Lines with a square at the top represent single breakends, and lines with arrowheads mark insertions.

**Supplementary Figure 27.**
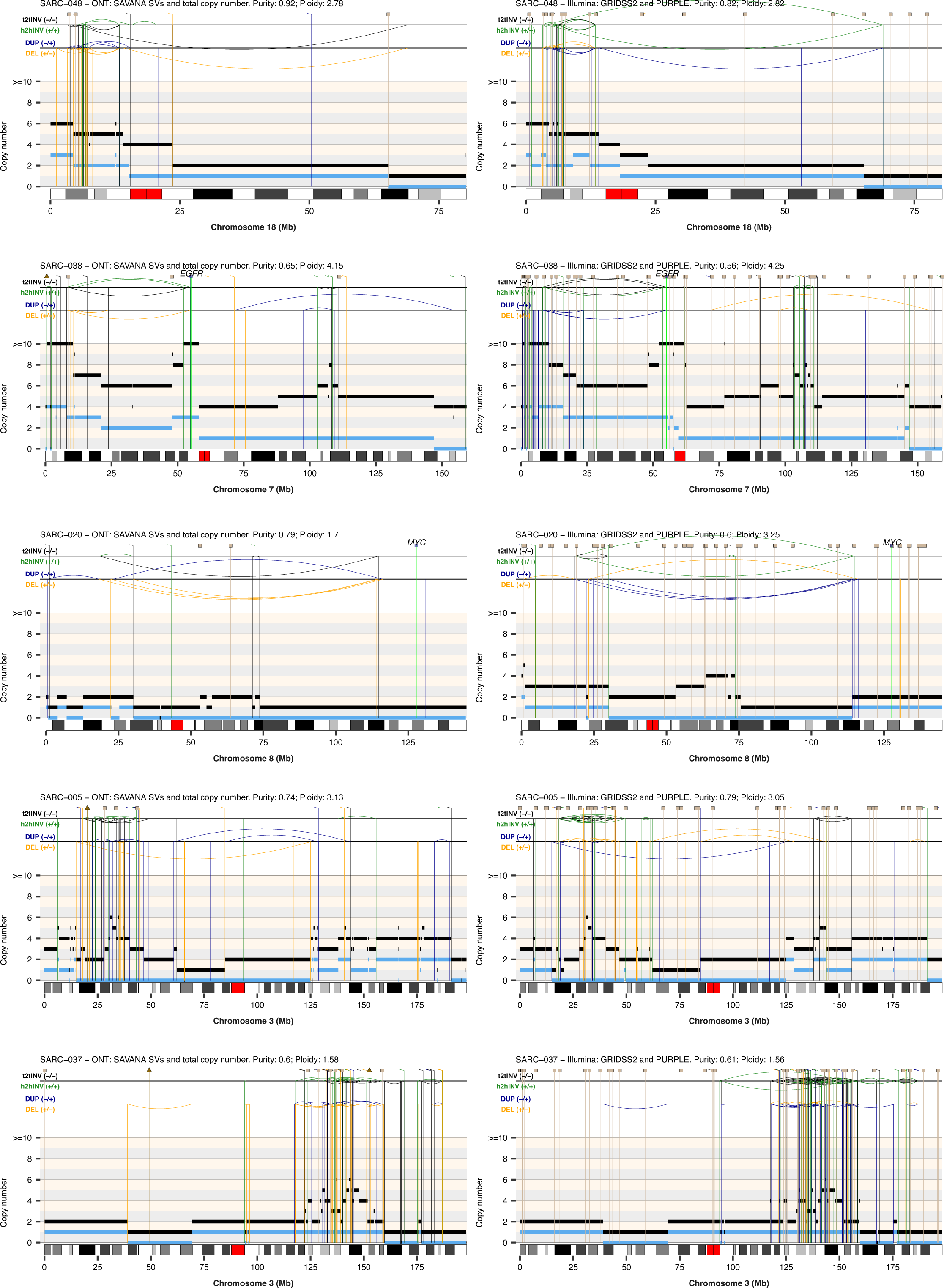
Somatic rearrangement and copy number profiles calculated using short-read data analyzed with GRIDSS and PURPLE against long-read data analyzed using SAVANA. (**Left**) Somatic SVs and SCNAs detected in matched long-read nanopore whole-genome sequencing data using SAVANA. (**Right**) Somatic SVs and copy number profiles detected using GRIDSS2 and PURPLE in whole-genome short-read sequencing data. The total and minor allele copy-number data are represented in black and blue, respectively. DEL, deletion-like rearrangement; DUP, duplication-like rearrangement; h2hINV, head-to-head inversion; t2tINV, tail-to-tail inversion. Lines with a square at the top represent single breakends, and lines with arrowheads mark insertions.

**Supplementary Figure 28.**
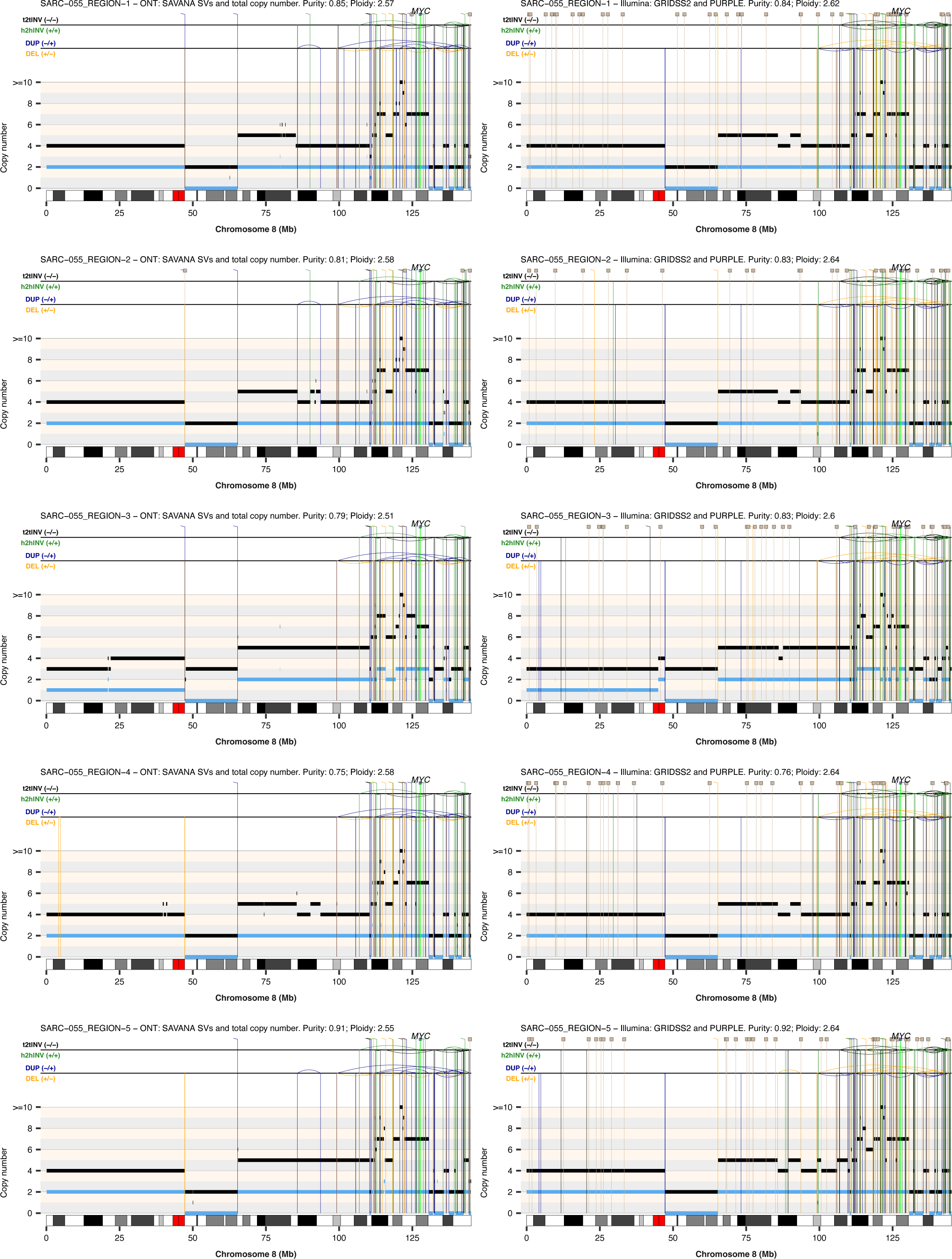
Somatic rearrangement and copy number profiles calculated using short-read data analyzed with GRIDSS and PURPLE against long-read data analyzed using SAVANA. (**Left**) Somatic SVs and SCNAs detected in matched long-read nanopore whole-genome sequencing data using SAVANA. (**Right**) Somatic SVs and copy number profiles detected using GRIDSS2 and PURPLE in whole-genome short-read sequencing data. The total and minor allele copy-number data are represented in black and blue, respectively. DEL, deletion-like rearrangement; DUP, duplication-like rearrangement; h2hINV, head-to-head inversion; t2tINV, tail-to-tail inversion. Lines with a square at the top represent single breakends, and lines with arrowheads mark insertions.

**Supplementary Figure 29.**
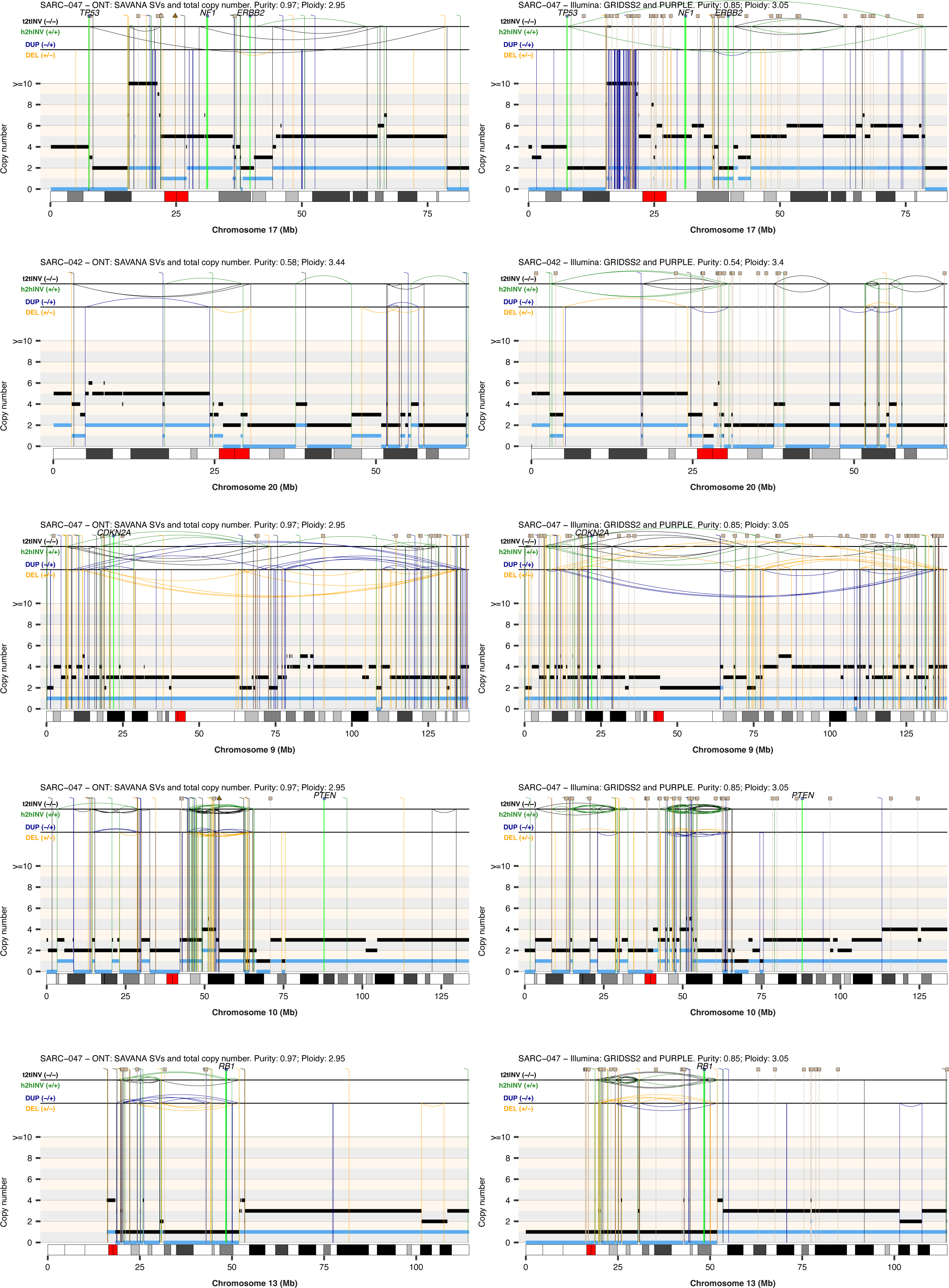
**Somatic rearrangement and copy number profiles calculated using short-read data analyzed with GRIDSS and PURPLE against long-read data analyzed using SAVANA.**(**Left**) Somatic SVs and SCNAs detected in matched long-read nanopore whole-genome sequencing data using SAVANA. (**Right**) Somatic SVs and copy number profiles detected using GRIDSS2 and PURPLE in whole-genome short-read sequencing data. The total and minor allele copy-number data are represented in black and blue, respectively. DEL, deletion-like rearrangement; DUP, duplication-like rearrangement; h2hINV, head-to-head inversion; t2tINV, tail-to-tail inversion. Lines with a square at the top represent single breakends, and lines with arrowheads mark insertions.

**Supplementary Figure 30.**
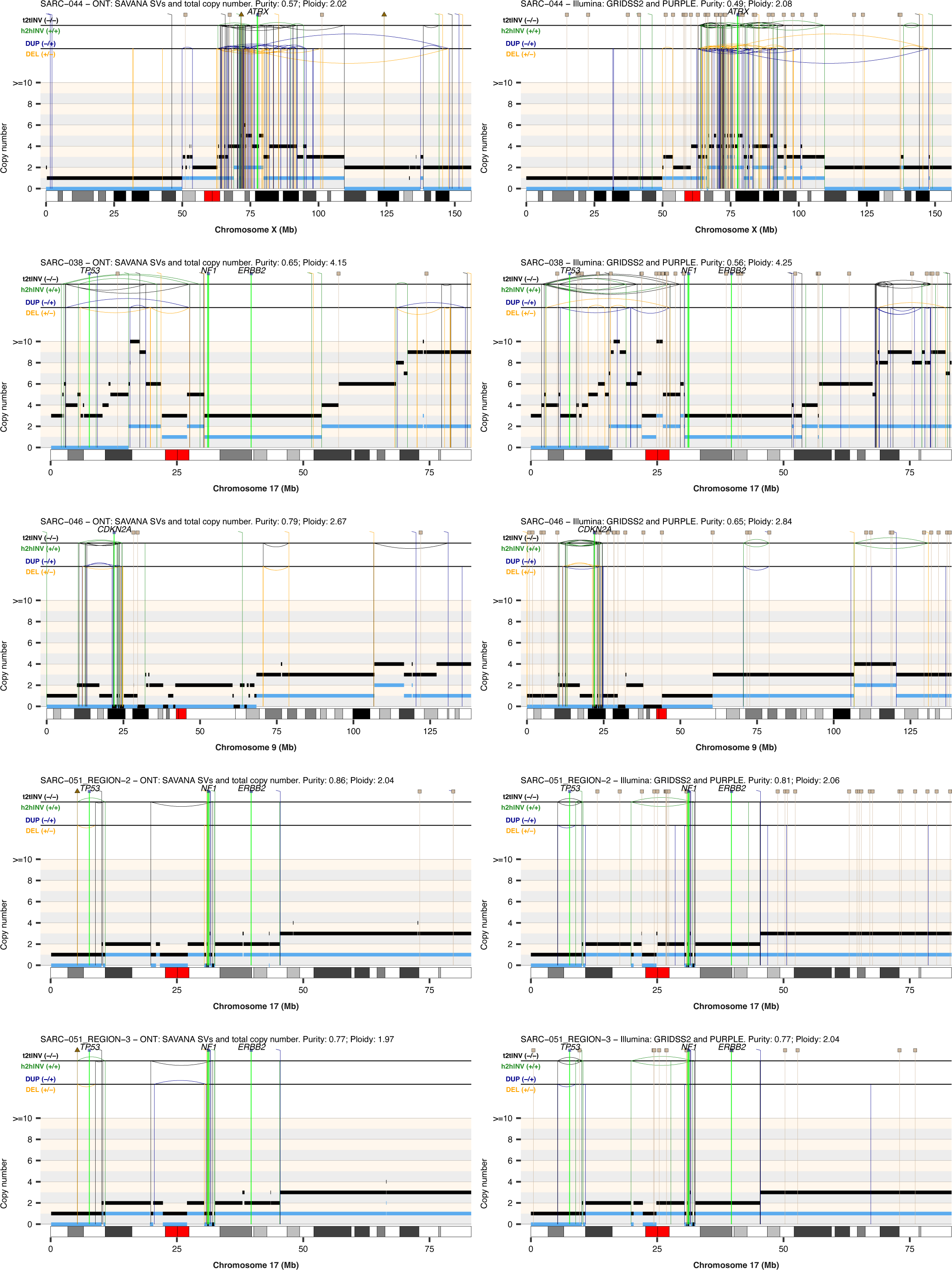
Somatic rearrangement and copy number profiles calculated using short-read data analyzed with GRIDSS and PURPLE against long-read data analyzed using SAVANA. (**Left**) Somatic SVs and SCNAs detected in matched long-read nanopore whole-genome sequencing data using SAVANA. (**Right**) Somatic SVs and copy number profiles detected using GRIDSS2 and PURPLE in whole-genome short-read sequencing data. The total and minor allele copy-number data are represented in black and blue, respectively. DEL, deletion-like rearrangement; DUP, duplication-like rearrangement; h2hINV, head-to-head inversion; t2tINV, tail-to-tail inversion. Lines with a square at the top represent single breakends, and lines with arrowheads mark insertions.

**Supplementary Figure 31.**
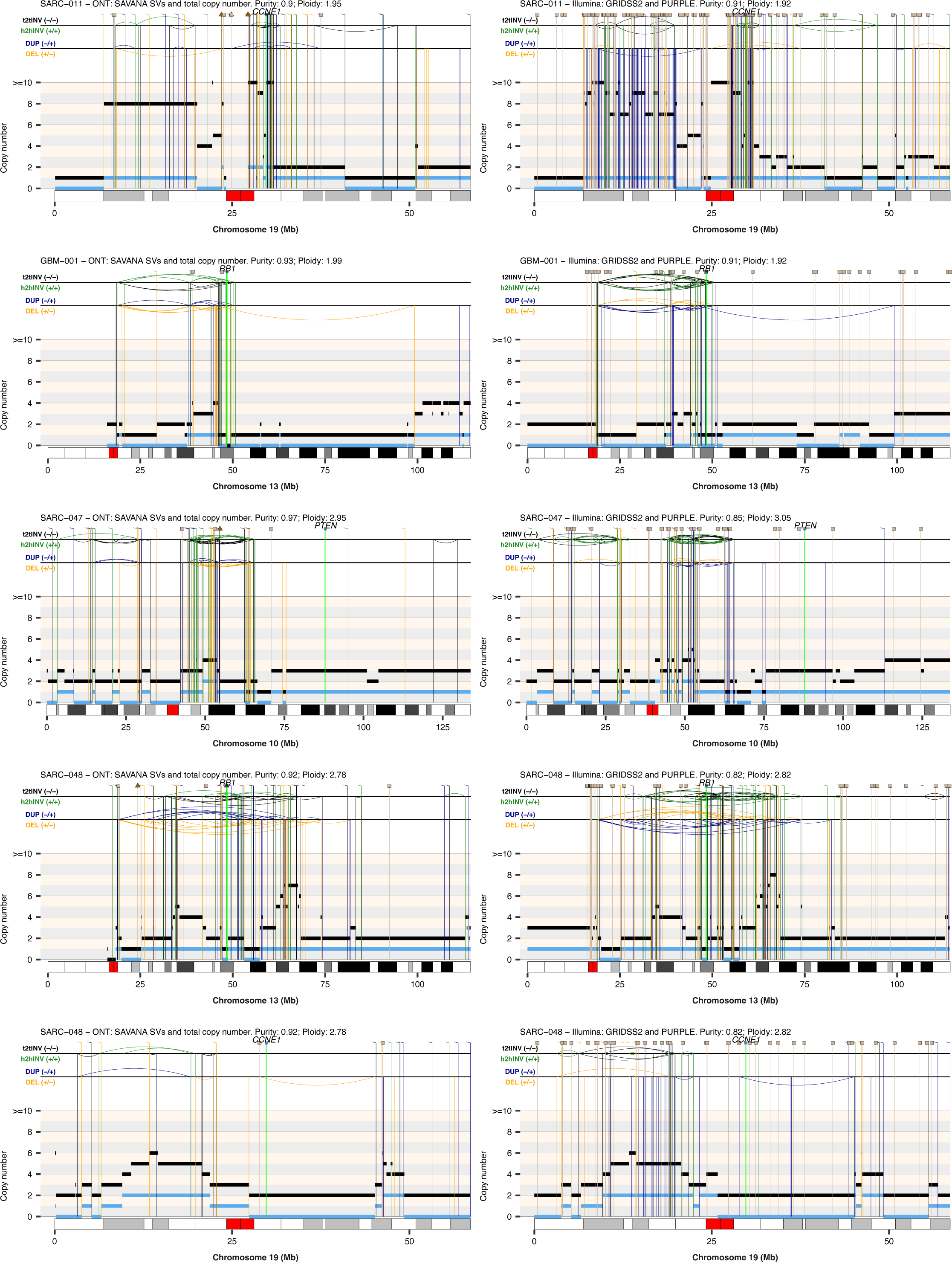
Somatic rearrangement and copy number profiles calculated using short-read data analyzed with GRIDSS and PURPLE against long-read data analyzed using SAVANA. (**Left**) Somatic SVs and SCNAs detected in matched long-read nanopore whole-genome sequencing data using SAVANA. (**Right**) Somatic SVs and copy number profiles detected using GRIDSS2 and PURPLE in whole-genome short-read sequencing data. The total and minor allele copy-number data are represented in black and blue, respectively. DEL, deletion-like rearrangement; DUP, duplication-like rearrangement; h2hINV, head-to-head inversion; t2tINV, tail-to-tail inversion. Lines with a square at the top represent single breakends, and lines with arrowheads mark insertions.

**Supplementary Figure 32.**
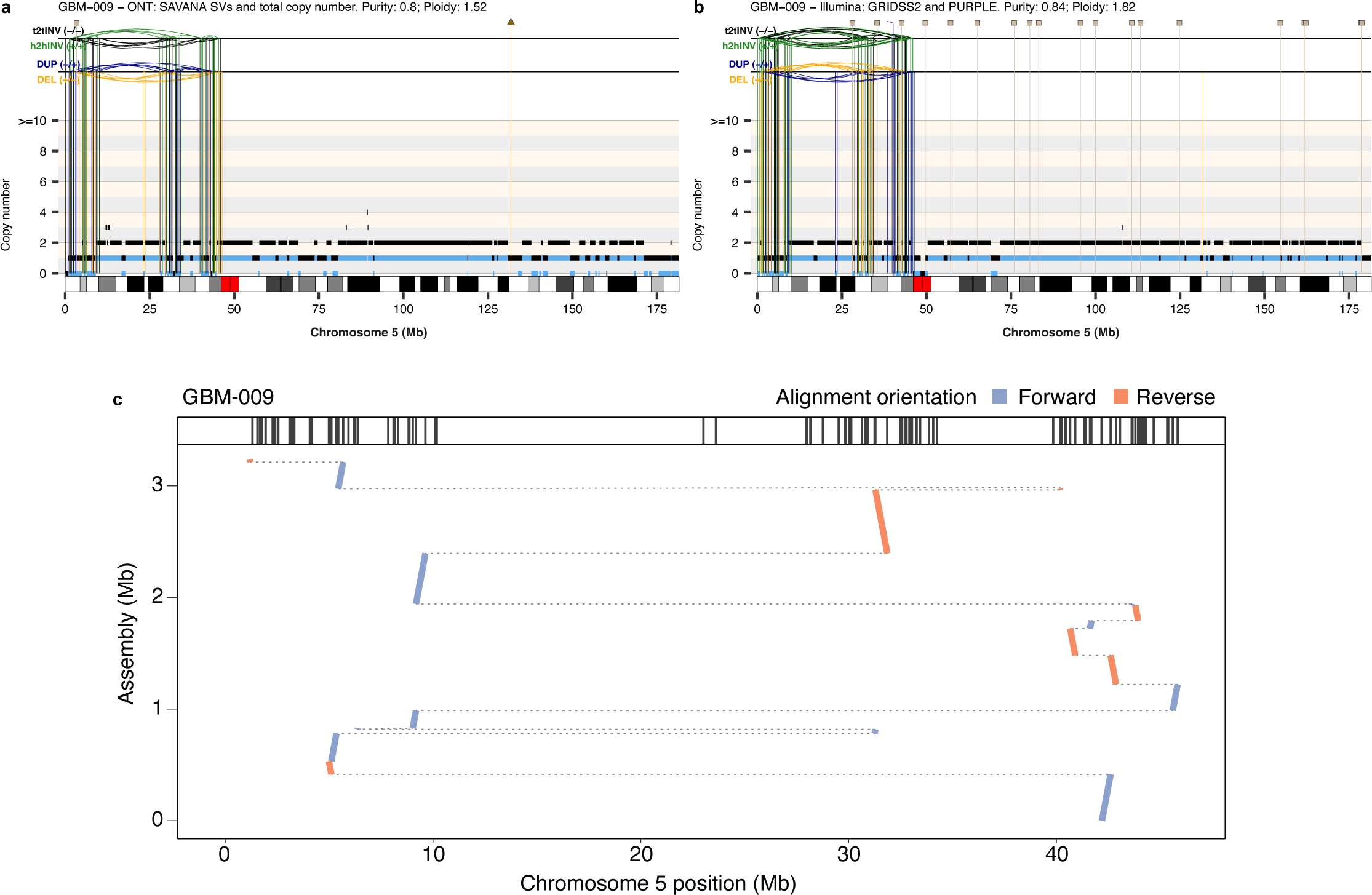
Application of haplotype-resolved somatic SV analysis using SAVANA to the assembly of derivative chromosomes generated by complex genomic rearrangements. Somatic SV and SCNA profile for chromosome 5 (**a-b**) involving a complex genomic rearrangement in tumour GBM-009. The total and minor allele copy-number data are represented in black and blue, respectively. DEL, deletion-like rearrangement; DUP, duplication-like rearrangement; h2hINV, head-to-head inversion; t2tINV, tail-to-tail inversion. Lines with a square at the top represent single breakends, and lines with arrowheads mark insertions. (**c**) Assembly of the complex genomic rearrangement shown in **a-b** using haplotype-resolved SVs detected by SAVANA at. The plot illustrates the alignment of the assembly to chromosome 5 of the human reference genome, revealing multiple connected SVs across the chromosome. The vertical lines in the top panel represent the coordinates of the somatic breakpoints detected by SAVANA.

